# Birth, cell fate and behavior of progenitors at the origin of the cardiac mitral valve

**DOI:** 10.1101/2022.08.06.503022

**Authors:** Batoul Farhat, Ignacio Bordeu, Bernd Jagla, Hugo Blanc, Karine Loulier, Benjamin D. Simons, Emmanuel Beaurepaire, Jean Livet, Michel Pucéat

**Affiliations:** INSERM U1251/Aix-Marseille University, Marseille, France; Department of Applied Mathematics and Theoretical Physics, Centre for Mathematical Sciences, Wilberforce Road, Cambridge CB3 0WA, UK; Wellcome Trust/Cancer Research UK Gurdon Institute, University of Cambridge, Tennis Court Road, Cambridge CB2 1QN, UK; Departamento de Física, Facultad de Ciencias Físicas y Matemáticas, Universidad de Chile, Santiago, Chile; Pasteur Institute UtechS CB & Hub de Bioinformatique et Biostatistiques, C3BI, Paris; Laboratory for Optics and Biosciences, Ecole polytechnique, CNRS, INSERM, IP Paris, Palaiseau, France; Sorbonne Université, INSERM, CNRS, Institut de la Vision, Paris, France; Wellcome Trust-Medical Research Council Stem Cell Institute, Jeffrey Cheah Biomedical Centre, University of Cambridge, Cambridge CB2 A0W, UK

## Abstract

Congenital heart malformations often include mitral valve defects which remain largely unexplained. During embryogenesis, a restricted population of endocardial cells within the atrioventricular canal (AVC) undergoes endothelial to mesenchymal transition (EndMT) to give rise to mitral valvular cells. However, the identity, fate decisions of these progenitors as well as the distribution of their derivatives in valve leaflets remain unknown.

Here, we use scRNA-seq of genetically labeled mouse AVC endocardial cells and of micro-dissected embryonic and postnatal mitral valves to characterize the developmental road. We uncovered the genetic, cell signaling and metabolic processes underlying specification of the progenitors and how they contribute to subtypes of endothelial and interstitial embryonic and postnatal valvular cells. Using clonal genetic tracing with multicolor reporter, we describe specific modes of growth of endocardial cell-derived clones which build up in a proper manner functional valve leaflets.

Our data reveal how both genetic and metabolic specification mechanisms specifically drive the fate of a subset of endocardial cells toward valve progenitors and their distinct clonal contribution to the formation of the valve.

## INTRODUCTION

During evolution, as organisms adapted to new environments, the heart has grown in size and acquired a complex structure, ranging from a simple contractile vessel in Amphoxius to a four chambers organ in mammals. These changes have taken place in concert with the development of valves that ensure unidirectional blood flow.

Malformation of valves accounts for around 10% of congenital heart diseases (CHD) (Hoffman and Kaplan, 2002) and can originate from early stage of development such as endocardial cushion defects or atrioventricular septal defects, which are common congenital heart malformations accounting for 7.4% of all CHDs (Digilio et al., 2019).

Mutations of a few genes have been reported as responsible for valves defects. These include genes encoding cytoskeleton proteins fibrillin and filamin A (Dietz et al., 1991; Kyndt et al., 2007), TGFβ receptors and the downstream signaling smad pathway(Neptune et al., 2003) and, more recently, Dachsous, potentially LMCD1, tensin and DZIP (Durst et al., 2015; Toomer et al., 2019). Altogether, mutations of these genes account for less than 5% of genetic valvulopathies. This suggests that, besides mutation of genes, other patho-biological processes such as defects in early cell fate and lineage specification, in metabolic pathways and/or in cell migration and cell distribution within the leaflets might be responsible for valve diseases.

Valve development is a complex biological process involving several cell lineages first specified and determined by specific gene transcriptional changes, influenced by many extrinsic factors and cellular events including growth factors, cell-cell and cell-matrix interactions and mechanical cues (Luna-Zurita et al., 2010). Valves form as early as embryonic stage E9.0-E10 in the mouse embryo and develop as cushions in two discrete cardiac regions, the atrioventricular canal (AVC) and the outflow tract (OFT). The AVC valves comprise the mitral and tricuspid valves, while the OFT or semilunar valves include the aortic and pulmonary valves(Combs and Yutzey, 2009). The mitral valve is the most affected valve in cardiac congenital diseases.

Valve formation commences with the delamination of endocardial cells that become separated from the myocardium by a cardiac jelly. This event is followed by endocardial cell transition toward a mesenchymal phenotype. Cell proliferation within an extracellular matrix forms the two cushions. The endothelial-to-mesenchymal transition (endMT) is triggered by a series of morphogenetic factors with still unknown specific or time-dependent role. Signaling pathways including TGFβ/BMP-smad, Wnt/TCF, and Notch4 secreted by the myocardium as well as NFATc mediated pathways (Zhou et al., 2005) likely governed by a set of specific genes and transcriptional pathways participate in the EndMT processes. The most intriguing feature of this biological process is that it is restricted within a set of a few endocardial cells. Besides the signaling pathways, little is known about the biological processes and transcriptional networks required for the cell to initiate local EndMT. Furthermore, the specific transcriptomic signature or biological properties that confer cells a competency for EndMT are still unknown.

Afterwards, a series of morphogenetic events allows the cushion cells to further differentiate into specific cell types (e.g., fibroblast, smooth muscle cells, chondrogenic cells) that proliferate and expand to build up the valve leaflets. It remains to be known how early lineage restriction occurs and how do cells expand, move and distribute within the valve leaflet (Combs and Yutzey, 2009).

To address these questions, we used a single cell-RNA sequencing approach combined with clonal analysis with multicolor reporters (Figure 1A) and uncovered the origin, the metabolic and transcriptomic signatures of valvular endocardial cells, and characterized the behavior of the valvular endothelial (VECs) and interstitial (VICs) cells during formation of mitral valve leaflets.

**Figure 1:**
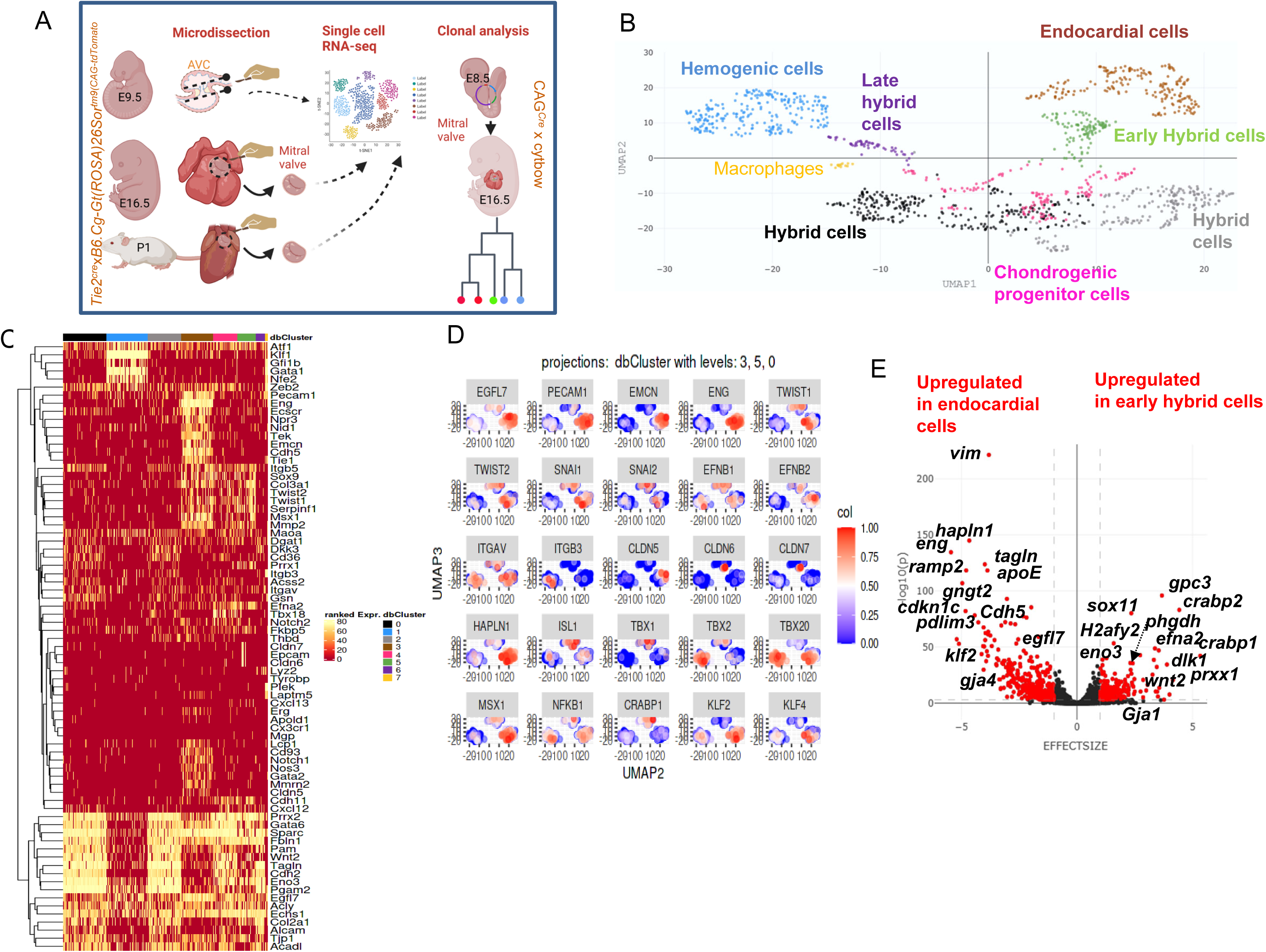
A: Experimental approaches B,C,D: single cell transcriptomic of E9.5 AVC cells. AVC was dissected from E9.5 embryonic hearts from a *Tie2^cre^*x*B6.Cg- Gt(ROSA)26Sor^tm9(CAG-tdTomato)^* mice breeding. Enzymatically dissociated cells were FACS- sorted and used for single cell-RNA seq. (B) UMAP showing the different AVC cell types (C) heatmap (D and E) panel plot and violin plot of differential gene expression in specific clusters 3 (endocardial cells) and 5 (early hybrid cells)

Our results show that endocardial cells constitute a heterogeneous cell population within the AVC. A subset of these cells which features a unique gene expression profile and which turns on specific and partially cilia-mediated metabolism pathways, are able to acquire EndMT competency and commit toward a specific valvular cell type. Based on our clonal and histological analyses, we estimate the number of self-renewing progenitors required to building a valve leaflet, and we observed multiple behaviors of VICs within the leaflet allowing for the formation of a functional valve.

## RESULTS

### Single-cell transcriptomics reveals cell specification in the atrioventricular canal

To interrogate the endocardial cell heterogeneity within the AVC, we first used a single cell-RNA sequencing approach. The AVC was carefully dissected out from 60 (E9.5) embryos (19-22 somites) from B6.Cg-Gt(ROSA)26Sortm9(CAG-tdTomato)Hze/J) females bred with Tie2*^Cre^* males (Figure 1A).

Cells were enzymatically dissociated, FACS sorted, and loaded into the chromium (*10X genomics*). 1500 cells were sequenced and 1200 which passed the quality control test used in the analysis. The algorithm for dimension reduction-generated Uniform Manifold Approximation and Projection (UMAP) derived plot (Figure 1B) revealed heterogeneity of the whole cell population. The AVC cell population featured an average of 5530 genes/cell and comprised one population of endocardial cells, two populations of hybrid cells undergoing EndMT, a subpopulation of late hybrid /mesenchymal cells as well as chondro-osteogenic progenitors and hemogenic cells. A small cluster of macrophages was also observed (Figure 1B).

The heatmap (Figure 1C) further revealed a broad spectrum of gene expression expected to mark future VIC and VEC populations. More specifically, we found one subpopulation of *eng^+^, pecam^+^, egfl7^+^, cdh5^+^, npr3^+^, tek^+^, tie1^+^, emcn^+^* endocardial cells (cluster 3), a subpopulation of *zeb2^+^, prrx1^+^, twist1,2^+^, msx1^+^, cldn7^+^, epcam^+^, efna2^+^* cells starting the transition from endothelial to mesenchymal fate (early hybrid, cluster 5), two subpopulations^-^ of high *tagln^+^, col2a1^+^ and twist1,2^-^* hybrid cells (clusters 0 and 2), and a subpopulation of *tagln^+^, col2a1^+^, fbln1^+^, and alcam+* late hybrid cells transitioning toward mesenchymal cells. We also observed subpopulations of *cdh11^+^, sox9^+^, sparc^+^, Tbx18+, col3a1^+^, cthrc1^+^, Gata6^+^, itgb5^+^, cxcl12^+^, maoa^+^* chondro-osteoblastic progenitor cells (cluster 4), of *Gata1*^+^, *nfe2*^+^, *klf1^+^*, *gfli1b*^+^ hemogenic cells (cluster 1) and a small cluster (cluster 7) of *Plek^+^, tyrobp^+^, Gata1*^+^, *nef2^+^, fcer1g^+^* macrophages. The endocardial cells of the AVC are thus already committed to specific valvular cell populations.

### Transcriptomics-driven EndMT

EndMT is the first step of valvulogenesis. We thus first addressed the question of the EndMT competency of these cells. Epithelial-to-mesenchymal transition (EMT) is no longer understood as a binary process but has been described as a continuum transition (Ognjenovic et al., 2020). EndMT is likely to follow such a step-by-step transition from endocardial to mesenchymal cell phenotype. We first interrogated a few genes that we surmised to play a role in EMT. The panel plots in Figure 2D and the violin plot in Figure 3E show only endocardial cells (marked by *pecam1, eng, cdh5* and *egfl7*) and early hybrid cells (marked by *twist1, twist2*). The plots reveal that only part of endocardial cells expressed *twist1, twist2,* together with *snai1,2*, an early marker of EMT. Interestingly these cells also expressed *klf2, klf4* and *msx1*, known to play a role in EndMT (Chen et al., 2008; Laurent et al., 2017). These cells lost *claudin 5*, an adherens junction gene together with the endocardial genes such as *cdh5*, or *eng* and the cell cycle inhibitor p57 (*cdkn1c*) while some early hybrid cells gained *claudin 6* and *7*, more reminiscent of migrating mesenchymal cells (Figure 1D). Expression of epithelial Cx37 (*gja4*) in endocardial cells switched to expression of Cx43 (*gja1*) a connexin involved in cell invasion in early hybrid cells (Figure 1E).

**Figure 2:**
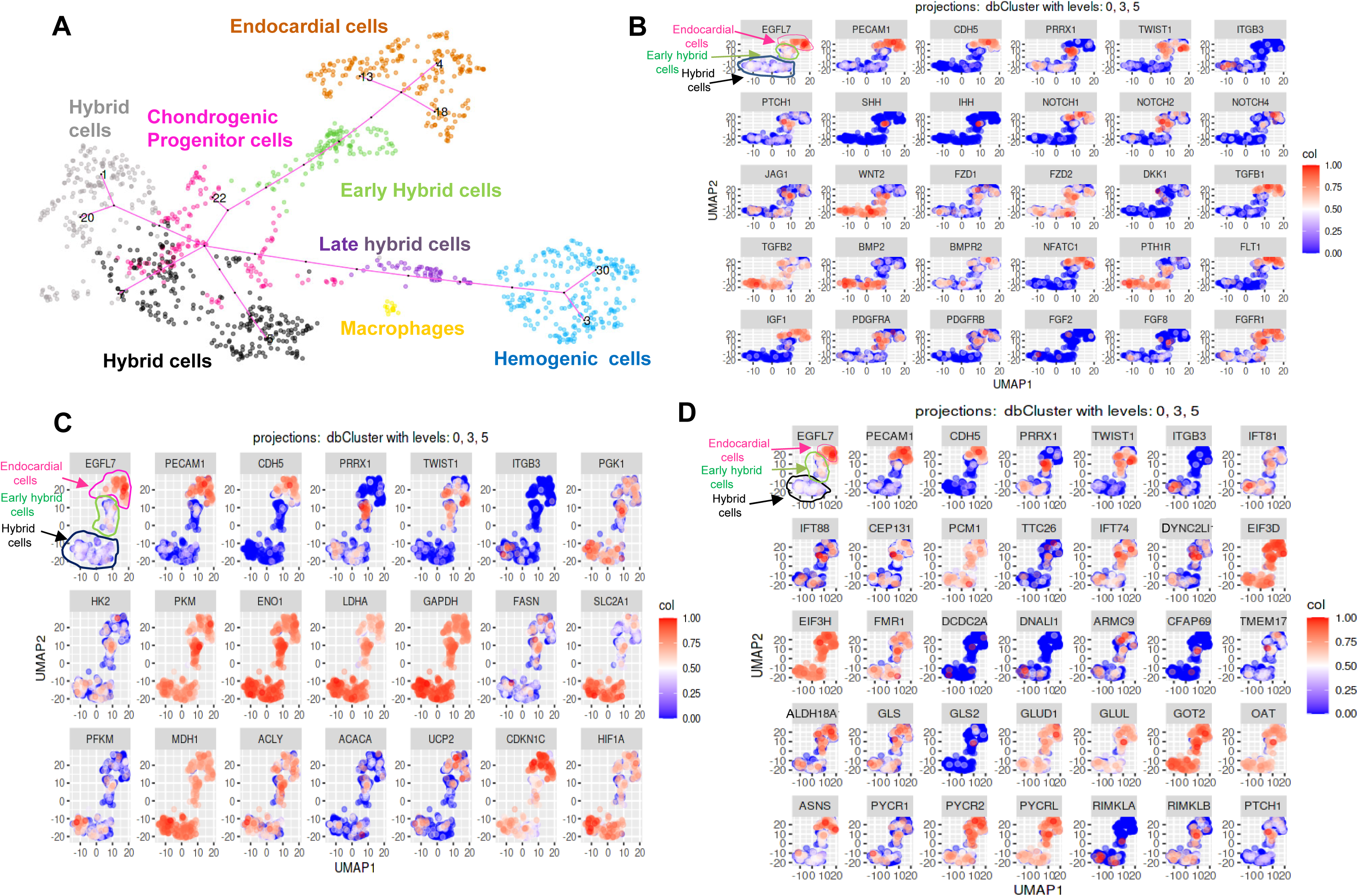
Trajectory inference analysis and signaling and metabolic pathways underlying EMT of endocardial cells within AVC. (A) elpigraph inference trajectory. (B) panel plots revealing expression of genes specifically involved in signaling pathways.(C) panel plots of genes involved in Warburg effect in cluster 3, 5 and 0 (endocardial cells (bottom left cluster 3), early (top cluster 5) and hybrid cells, (bottom right cluster 0) (D) panel plots of genes involved in Glutamine metabolism and cilia formation in cluster 3, 5 and 0 (endocardial cells (top cluster 3), early middle cluster 5) and hybrid cells, (bottom cluster 0)

**Figure 3:**
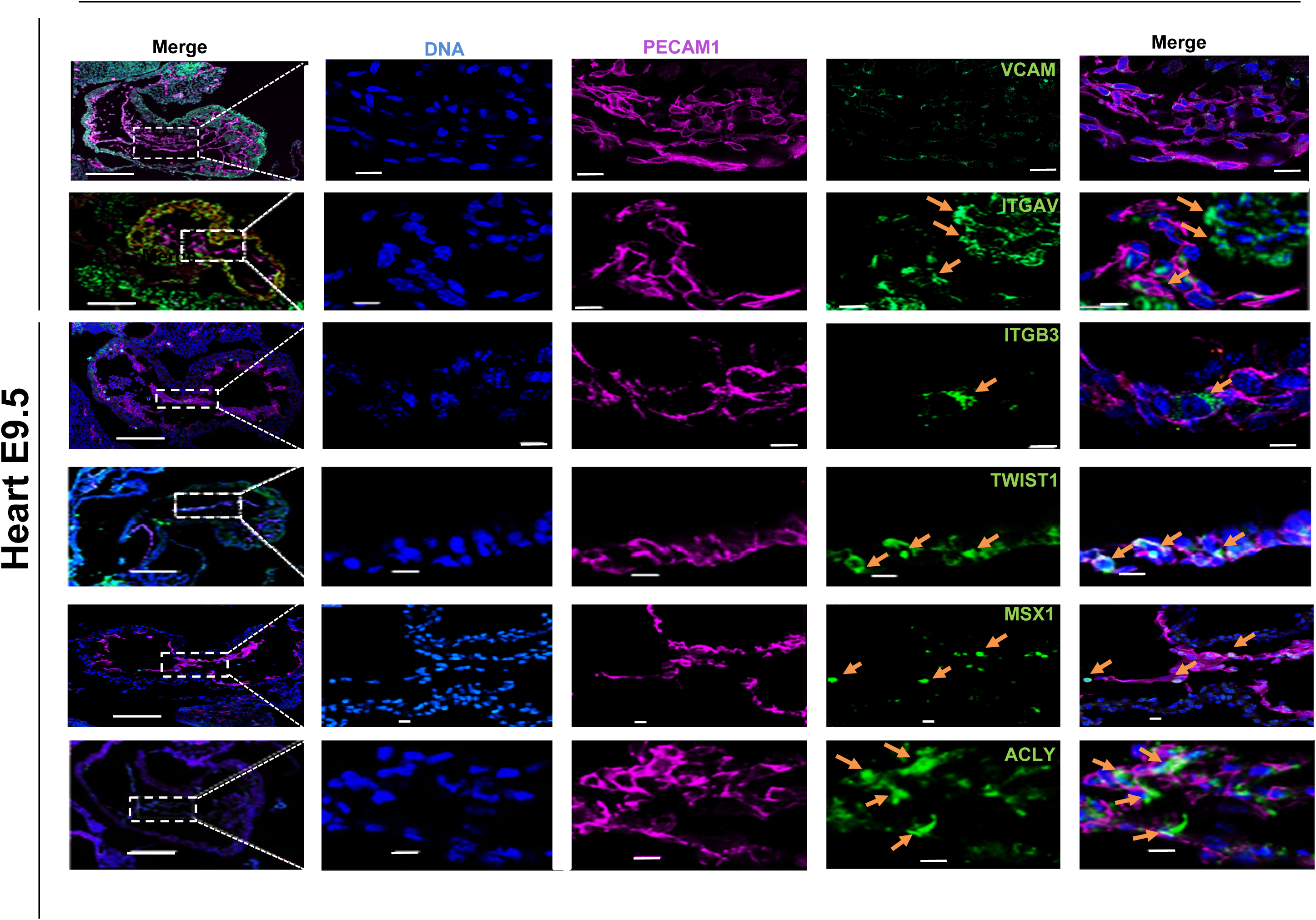
immunofluorescence of proteins expressed in AVC cells in E9.5 mouse heart. The left panels show low magnifications of sections of whole heart. The square marks the AVC only imaged in the other panels. Positive cells for the marker of interest are pointed by orange arrows. The scale bars in the left column is 100 μm and the others are 10 μm

A full analysis of differential gene expression between cluster 3 (endocardial cells) and 5 (early hybrid cells) can be found in supplemental data 1.

To further dissect this transition, we used an inference trajectory analysis. Interestingly, this bioinformatics approach suggested that two subpopulations of endocardial cells gave rise first to early *twist1^+^*hybrid cells. The cells then progressed towards hybrid mesenchymal cells still retaining expression of genes reminiscent of endothelial cells. Three branches emerged from these cells that are engaged into the process of EMT. They gave rise to either chondro-osteogenic cells, remained as hybrid cells, or moved towards late hybrid and hemogenic cells (Figure 2A). Macrophages remained slightly detached from the trajectory while remaining in the hemogenic cells trajectory.

### Signaling pathways –driven EndMT

More specifically, we looked at signaling pathways that could be involved in EndMT (Figure 2B). BMP2 was expressed only in hybrid cells (cluster 0) while TGF*β*1 was found in in endocardial cells (cluster 3) but not in hybrid cells. The VEGFR *flit1* was expressed only in endocardial cells. The *notch2 and SHH* and *IHH* ligands and their respective *Jag1* and *Ptch1* receptor genes were expressed mostly in early hybrid cells but not in endocardial cells*. IGF1* was also mostly expressed in endocardial cells. We also observed the expression of the parathyroid hormone receptor (*pth1R)* whose expression was switched on in early hybrid cells and upregulated in hybrid cells. FGF receptor 1 (*FGFR1*) was expressed in endocardial and early hybrid cells and decreased its expression in hybrid cells. Expression of *FGF8* but not of *FGF2* was mostly turned on in hybrid cells. Wnt2 expression was turned on in early hybrid and hybrid cells. PDGFRα, a receptor known to be present on the primary cilium, during valve development (Moore et al., 2021) was expressed like PDGFRβ, in endocardial and early hybrid cells (Figure 2B). Collectively our data point out a time-dependent role of growth factors during the EndMT process.

### Metabolism-driven EndMT

Interestingly, genes encoding key glycolysis enzymes such as hexokinase 2 (*hk2),* glyceraldehyde 3-phosphate dehydrogenase (*gapdh* phosphoglycerate kinase (*PGK1*), enolase 1 and 3 (*eno1,3),* phosphofructokinase (*pfkm*), and lactate dehydrogenase-A (*ldhA*) as well as the glucose transporter 1 (*Slc2A*) were expressed in some endocardial cells and all along EndMT. Altogether these data suggest that cells turned on the Warburg effect. The malate dehydrogenase (mdh1) the acetyl-CoA Carboxylase Alpha (*acaca*), the acetyl Co-A transferase (*acly*), the acetyl-CoA decarboxylase (*acss2*), the fatty acid synthetase (*fasn*), all involved in the Warburg effect were also expressed in combination with p57 (*cdknc1*) and *hif1a* (Courtnay et al., 2015) (Figure 2C) all along the EndMT process in restricted cell populations and in a few endocardial cells (Figure 2C). Their expression increased in early hybrid *twist1^+^*cells, and to a low extent in late hybrid cells (Figure 2C).

Together with the Warburg effect, aerobic glycolysis is often associated with the glutamine metabolism such as in cancer cells (Yoo et al., 2020). Furthermore, a recent report links the primary cilia known to play a role in the mitral valve morphogenesis as early as in the atrioventricular canal (Burns et al., 2019; Durst et al., 2015; Toomer et al., 2019) to the glutamine metabolism (Steidl et al., 2023). Thus, we interrogated a set of genes encoding enzymes involved this metabolic pathway as well as in cilia formation. Glutamine transporters genes *slc38a*, *slc1a5*, *slc38a2* were highly expressed in some endocardial cells and early hybrid cells while decreasing their expression in hybrid cells. A similar pattern of expression was observed for *pycr2*, *pycrL* encoding enzymes involved in proline synthesis. A*ldh18a2* gene encoding an aldehyde dehydrogenase gamma-glutamyl modifying protein followed the same profile of expression as *pycr2* and *pycrL* in endocardial and early hybrid cells. *Asns*, encoding the asparagine synthetase using glutamine as a subtrat was highly expressed in endocardial cells and slowly switched off in early and hybrid cells.

Genes encoding enzymes involved in glutamine synthesis (*gluL*) and in α−ketoglutarate synthesis and entry into the Krebs cycle and possibly mTORc1 activation, *gls*, *got2*, *glud1* as well as *idh1* were highly expressed in some endocardial cells and the expression was maintained along the EMT process.

*RimklA and rimklB (Ribosomal Modification Protein RimK Like Family Member B)*, genes encoding enzymes catalysing N-acetyl-L-aspartate-L-glutamate and citrate-L-glutamate ligases were observed mostly in early and hybrid cells. Altogether the profile of expression of these glutamine metabolism genes suggests a cell requirement for high activity of this metabolism important both for non-essential amino acid synthesis and mTor pathway.

The genes involved in cilia formation including ift81, ift88, ift74, cep131, pcm1,ttc26, dync2Li1,Eif3D, Eif3H, fmr1, dcdc2A, dnali1, armc9, cfap69 and Tmem17 were expressed at high level in some endocardial cells (but dcdc2a and dnaLi1 and cfa69) and all along EndMT (Figure 2D). Most of them were co-expressed with the genes encoding the enzymes involved in the glutamine metabolism and asparagine synthesis (Figure 2D) Altogether, this profile of gene expression points to a key role in EndMT process of the Warburg effect and of cilia formation together with the glutamine metabolism.

### Protein expression in AVC

We next picked up proteins encoded by genes of interest in the EndMT process and carried out immunofluorescence experiments on sections of E9.5 embryonic heart. Vcam1 was barely or not expressed in endocardial Pecam+ cells and Twist1, a protein involved in early EMT, featured a salt and pepper staining pattern in the endocardium within the AVC as shown in the panel plot (Figure 3). TWIST was still located into the cell cytosol and TWIST+ cells also co-expressed Pecam1. MSX1, another marker of EndoMT turned on by Bmp2 signaling from the myocardium, was observed in delaminated cells migrating from the endocardium to the cushion. The fourth marker, the ATP-citrate lyase (ACLY), was expressed in some endocardial cells, as well as in cells migrating within the cushion still expressing pecam1, namely, early hybrid cells. Two other markers for late hybrid EndoMT and mesenchymal cells were validated: ITGAV was expressed in endocardial and post-EndoMT cells in the cushion, and ITGB3 was only expressed in rare delaminated endocardial cells. MSX1 was also expressed in very few cells of delaminated endocardium (Figure 3).

These observations demonstrate that an activation of glycolysis and the Hif1α dependent Warburg effect and glutamine metabolism are crucial events in the EndMT process. Furthermore a time-dependent consecutive activation of TGFβ1, then SHH, and Notch and finally PTH and Wnt pathways confer to endocardial cells a potential to undergo EndoMT.

### Transcriptomic profile of embryonic and postnatal mitral valve depicts early valvulogenesis

We collected embryos and neonates generated from B6.Cg-Gt(ROSA)26Sortm9(CAG-tdTomato)Hze/J) females bred with Tie2*^Cre^*males at E16.5 and P1, respectively. Hearts were dissected out, opened and the annulus of the mitral valve including the leaflets were carefully cut out, enzymatically dissociated, and pooled together from 92 embryos at the same stage of development (Figure 1A). The isolated cells were sorted by FACS and altogether 954 Tomato+ cells were then subjected to single-cell RNA sequencing and analysis.

The UMAP revealed that E16.5 valvular cells segregated between VEC expressing *pecam1, cdh5, and eng* and VICs expressing markers of fibroblasts, *postn and dcn* (Figure 5A). VECs distributed into two clusters including one with cells expressing *wt1, and not CD36*, a fatty acid transporter suggesting that these VECs belong to the parietal leaflet in which epicardial cells contribute to (Wessels et al., 2012) (Figure S2) and the other one with cells expressing *CD36* and not *wt1* (Figure 4A).

**Figure 4:**
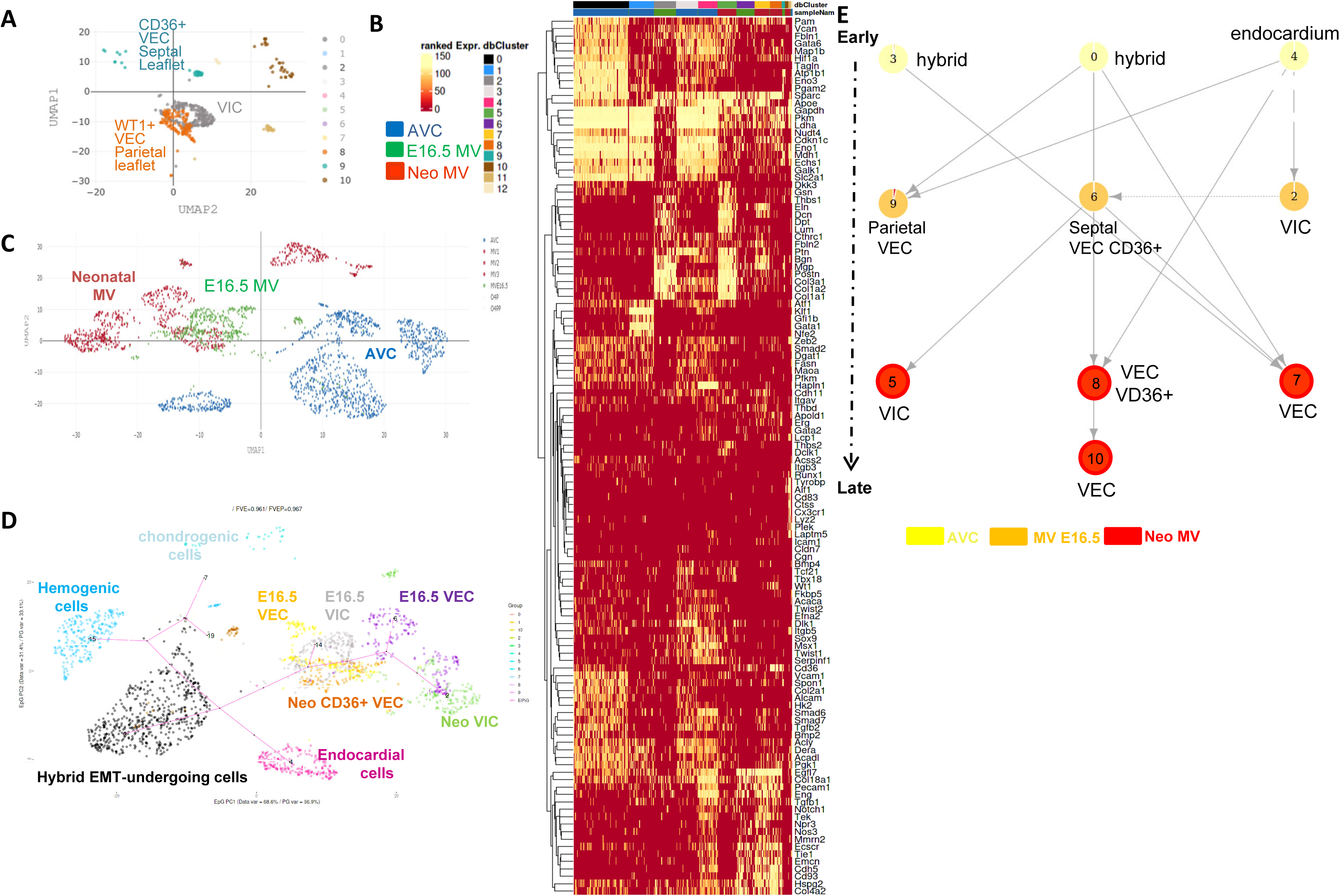
single cell transcriptomic of E16.5 and neonatal mitral valves (MV). Mitral valves were dissected from embryos and neonates from *Tie2^cre^*x*B6.Cg- Gt(ROSA)26Sor^tm9(CAG-tdTomato)^* breeding. Enzymatically dissociated cells were FACS-sorted and used for single cell-RNA seq. (A) UMAP of E16.5 MV cells, (B) heatmap of gene expression in AVC, E16.5 MV and neonatal (Neo) MV cells.(C) UMAP of all samples (AVC, E16.5 and neonatal mitral valve cells). (D) elpigraph inference trajectory including all datasets. (F) tempora inference trajectory of all samples

**Figure 5:**
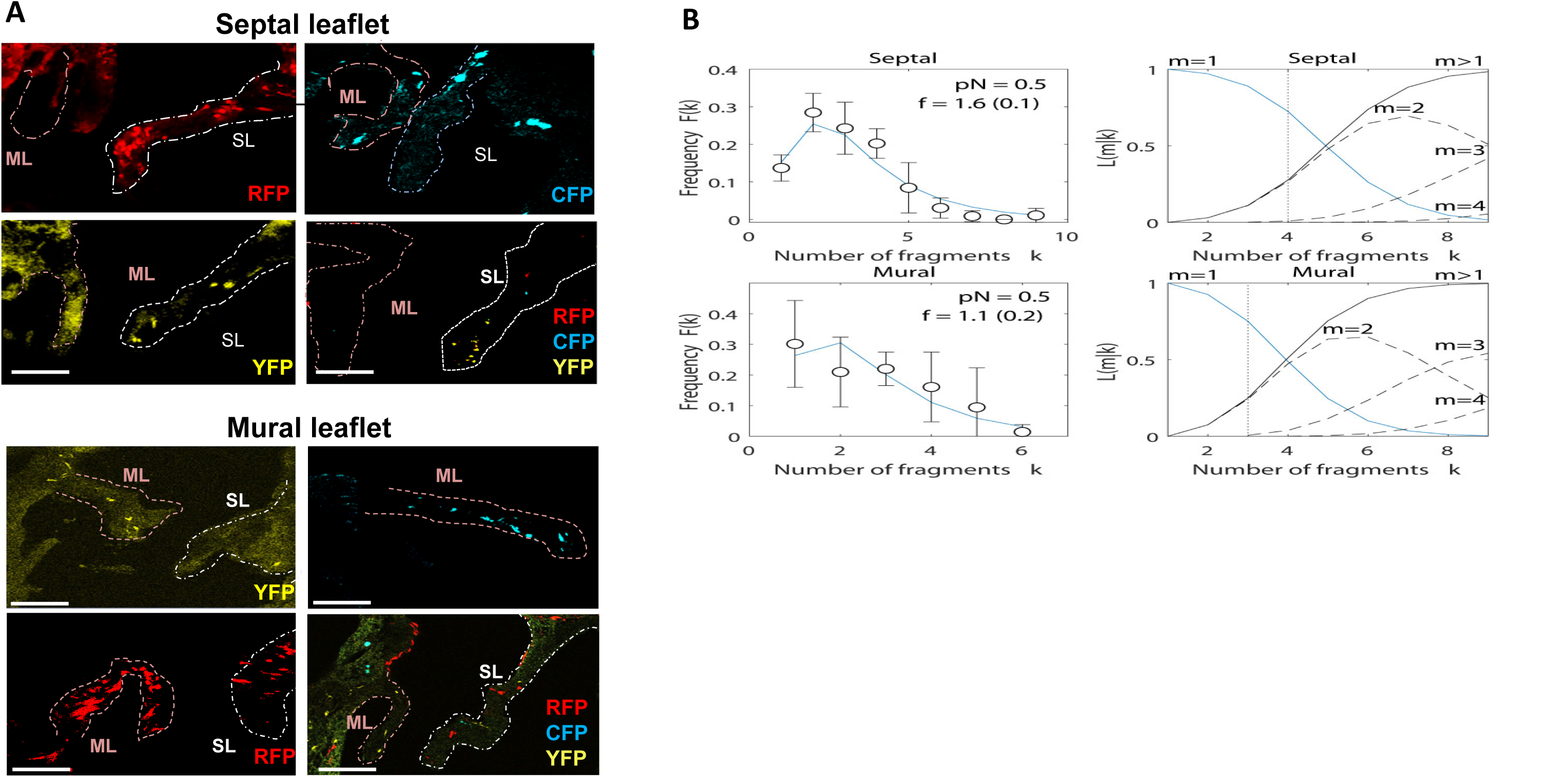
Cell clone distribution in embryonic mitral valve. (a) E16.5 mitral valve leaflets images with cell clones from CAG*^Cre^* x cytbow embryos (scale bar 50 μm) (b) Probability F(k) of observing a total of k fragments of any given color, and (right) conditional probability L(m|k) that m cell were induced given that we observe k fragments in the anchoring point for the (top) septal and (bottom) mural leaflets. Markers correspond to experimental measurements, error bars correspond to the standard deviation from the measurement of F(k) for the three different colors. The solid line was obtained from fitting Eq. (5) in methods for pn = 0.50 and free parameter f.

E16.5 valvular cells segregated from the neonatal valvular cells as well as AVC cells segregated from E16.5 and P1 valvular cells as shown in the heatmap and UMAP plot (Figure 5BC). The heatmap including AVC, E16.5 embryonic and neonatal mitral valve cells revealed the presence of numerous valvular genes (Figure 4B). *Egfl7, pecam1, cdh5, eng, ecscr, tie1, cd93, thbd, and emcn* were expressed in endocardial cells (AVC cluster 4), and VEC of E16.5 mitral valve (cluster 5,6) and of neonatal mitral valve (clusters 7 and 8). Interestingly some E16.5 VEC and neonatal VEC (cluster 7 and 8) expressed *CD36* and *fabp4*, two markers of fatty acid transport and metabolism.

VICs of both E16.5 and P1 embryonic and neonatal mice (clusters 2 and 6) expressed numerous markers of VIC of adult valve layers (spongiosa, fibrosa and atrialis). Indeed, we interrogated a list of genes specific of spongiosa *(dpt, postn, thbs2, eln, col3a1, col1a1), fibrosa (col1a2, thbs1, bgn, Dkk3)* or atrialis enriched in elastin *(eln).* All genes were expressed in most cells of both E16.5 and neonatal mitral valves in clusters 2 (E16.5) and 6 (neonatal) even in cells that did not cluster in a layer-specific manner. Immune cells, including macrophages which expressed *tyrobp, plek, ctss, laptm5, runx1, cd83, aif1, lyz2 and cx3cr1,* were found in cluster 10 (Figure 4B).

We then applied linear trajectory inference via pseudotime analysis (using Elpigraph) to track the steps of valvulogenesis (from day E9.5 until post-birth). The trajectory described the multistep process by which AVC cells at E9.5 transition towards valvular cells. This analysis revealed a subset of *pecam1+, eng+, cdh5+* endocardial cells that give rise to the endothelial layers (VEC) of both E16.5 and neonatal valves. The endocardial cells also contributed to subsets of VICs. One subset at E16.5 led to the VICs in neonatal valve. The endocardial cells also generated hemogenic cells and underwent EMT to give rise to chondrogenic cells (Figure 5D). We more specifically interrogated ECM-related genes in the three single-cell RNA-seq datasets (AVC, mitral valve at E16.5 and at neonatal stages). The heatmap in Figure S3 showed that clusters 1 and 4 (including only E16.5 and neonatal valvular cells) were enriched in collagen genes (*col1a1, col1a2, col3a1)*, reminiscent of the fibrosa layer. Clusters 5, 7 and 8 including E16.5 and neonatal mitral valvular cells were enriched in proteoglycans including perlecan (*hspg2)*, versican (*vcan),* lumican *(lum),* and elastin (*eln*), reminiscent of the spongiosa and atrialis layers. AVC cells from clusters 0, 2, 3 and 6 specifically expressed the galactose kinase (*galk1*), an enzyme required to synthetize proteoglycans, as well as the hyaluronan receptor CD44 and the hyaluronan synthase 2 (*has2*).

Next, we asked whether the Warburg effect was conserved after EndMT in VICs of E16.5 and neonatal mitral valves. Figure S3 shows that both VECs and VICs at both stages of development maintained a glycolytic metabolism and a Warburg effect. Figure S4 also reveals that the genes involved in cilia formation and glutamine metabolism were still on in E16.5 and neonatal mitral valves and more specifically in pecam1+ VECs.

We further used Tempora (Tran and Bader, 2020) another cell trajectory inference using biological pathways information. We used a minimum size of gene sets of 50 and a maximum to 300 to allow the algorithm to use the most defined biological pathways.

This trajectory first revealed that cells differentiate step by step at each stage of development to finally contribute to post-natal valve leaflets.

The endocardial cells (cluster 4) as well as hybrid cells (clusters 0) contributed directly to the E16.5 parietal leaflet VEC (cluster 9), E16.5 VIC (cluster 2) and to neonatal VEC (clusters 8,10). The septal leaflet at E16.5 received cells from VICs (cluster 2) originating from endocardial cells (cluster 4) as indicated by the elpigraph trajectory (Figure 4D) Interestingly, CD36+ septal E16.5 VEC from cluster 6 from neonatal valve contribute to both CD36+ and CD36 low neonatal VEC (cluster 8 and 7, respectively) of the valve as well as to neonatal VICs (cluster 5) (Figure 4E).

The cluster 3 of hybrid cells contributed to neonatal mitral valve VECs while endocardial cells contributed to neonatal CD36+ VECs (cluster 8).

### AVC-derived valvular cell clones expand and fragment along the leaflets’ axis

The embryonic cell lineage at the origin of endocardial cells is diverse, which could explain their transcriptomic heterogeneity and the competency to EndMT being only restricted to specific cell types. We therefore questioned how post-EMT cells further acquire their fate and distribute within the valve leaflets. To explore the behavior of valvular cells during the formation of the valve leaflets, we used an experimental approach that is complementary to the analysis of their unique transcriptomic profile. We set up a multicolor lineage tracing approach using CAG-*CreER^TM^* male bred with Cytbow females (Figure 1). The latter line expresses a modified Brainbow transgene that stochastically triggers either cyan, or yellow, or red fluorescent protein expression upon Cre/lox recombination. The recombinase activity was induced at E8.5 using a low dose of tamoxifen (20 μg/g mouse) which sparsely label pre-EMT progenitors of valvular cells (Figure S5 A,B). Embryos were first collected at E9.5 to check the number of labelled endocardial cells. Extended data Figure 3A shows the labelling of a single endocardial cell at the proposed tamoxifen dosage. Phosphohistone 3 (PH3) immunostaining was used to score the rate of cell division (i.e. DNA duplication) of AVC endocardial cells at E8.5. A low cell proliferation rate was observed at E9.5 (Figure S5B) implying that cell division right after recombination was limited.

We first monitored labeled cells in the mitral valve of E16.5 embryos via color two-photon microscopy (Mahou et al., 2012) of whole clarified hearts. Large volume 3D microscopic imaging revealed that labeled cells formed elongated clusters that distributed primarily along the long axis of the leaflet (Figure S6A). This morphology resulted in a high correlation between the 2D-projected area of cell clusters and their 3D distribution (Pearson’s r=0.91, p=0.01, Figure S6B), which allowed us to base our clonal analysis on tissue sections. (Figure A,B).

Within a septal or mural leaflet we could observe that putative clones of a given color where fragmented into smaller clusters of cells (Figure 5A). To infer whether a group of clusters of cells belonged to a single clone, we followed an established biostatistical strategy (Wuidart et al., 2018) that allowed us to estimate the likelihood that a group of clusters was monoclonal (see Methods for details).

Applying this approach to the full clonal data, we first estimated both the overall induction frequency, pN (the frequency at which a single leaflet progenitor is induced, times the total number of leaflet precursors), and the clone fragmentation rate f (the average number of fragments per clone) (Figure 5B and Figure S7). By considering the combined data from both mural and septal leaflets, we estimated the overall induction frequency to be pN=0.5±0.3 (95% C.I.). By contrast, the fragmentation rate was estimated independently for each leaflet, giving f=1.6±0.1 for the septal leaflet, and a slightly lower rate of f=1.1±0.2 for the mural leaflet. From this, we found that, in the septal leaflet, groups of 4 or less cell clusters were likely to belong to a single clone while, in the mural leaflet, this threshold was reduced to 3 fragments (see Figure 6B and Methods for details). Based on this estimate, out of the 106 clone candidates, 83% (54 clones for the septal and 34 clones for the mural leaflet) satisfied the constraint and could be considered as clonal with statistical confidence.

**Figure 6:**
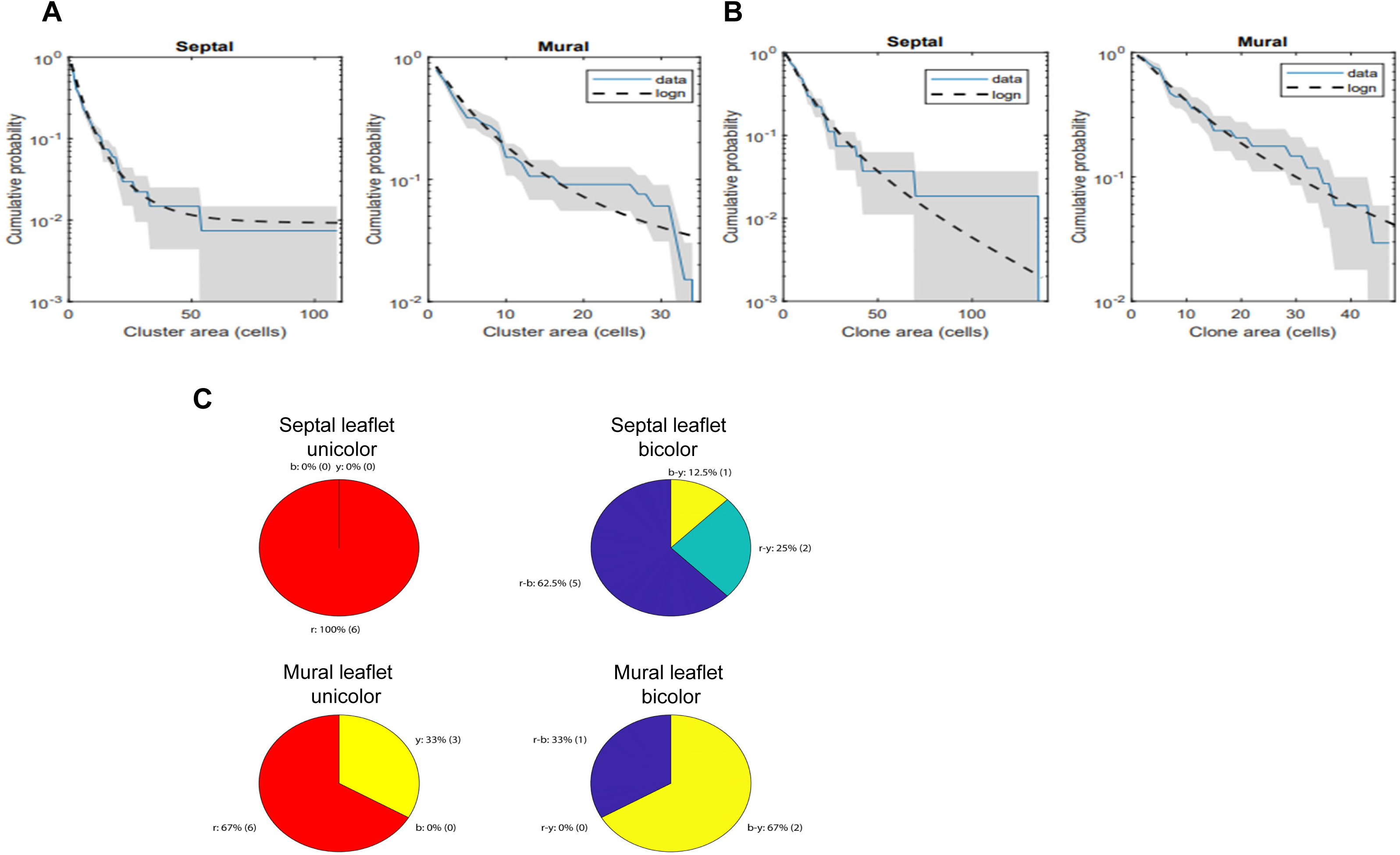
Distribution of clusters (A) and cell clone (B) sizes in both septal and mural leaflets of E16.5 mitral valve. (logn:log normal distribution) (C) observed color expression in both septal and mural leaflets compared with ramdom combinations

### Fragment and clone size distribution

Focusing on the clones identified by our statistical analysis, the distributions of cluster size within clones for both leaflets were found to fit well with a log-normal dependence (Figure 6A), with logarithmic mean µs = 1.3 (2) and variance σs = 0.9 (2) for the septal, and µm = 1.4 (2) and σm = 1.0 (2) for the mural leaflet. Indeed, such log-normal behavior has been shown to be a hallmark of clone fragmentation/coagulation processes (Bailey, 1964; Chabab et al., 2016; Redner, 1990), lending additional support to the assignment of clones from clusters. The fact that the size distribution of clusters for both leaflets was statistically indistinguishable suggested that both leaflets share a similar growth dynamic.

Similarly, the distribution of total clone sizes (comprising the sum of constituent clusters) could be fit well by a log-normal dependence (Figure 6B). A log-normal distribution of clone sizes could result from a cooperative growth process in which the stochastic proliferation of cells in the same neighborhood is positively correlated (i.e., clones arise as the “product of a random number of symmetric cell divisions before differentiation”). To challenge this hypothesis, we tested whether the sizes of clones of different color in the same leaflet were positively correlated (see Figure S8). From this, we found that indeed some clones sizes where positively correlated, while other others were not, suggesting a dependence on the spatial location of individual clones, and observation that was consistent with the suggested growth dynamics.

### Estimation of the number of progenitors of the mural and septal leaflets

We proceeded to estimate the number of progenitors contributing to the formation of the mural and septal leaflets. To this end, we estimated the average number of clones needed to cover the 2D-projected area of each of the leaflets. For the septal leaflet, we estimated that about 41±8 (SEM) dividing progenitors were required to cover the cross-section of the leaflet, while about 22±4 (SEM) progenitors were required in the case of the mural leaflet. Given the similarities observed in the clonal organization and estimation of number of progenitors in both leaflets, we questioned whether the pool of progenitors could be the same for both leaflets, so that there is a unique common pool of about 40 progenitors that give rise to both leaflets. In this spirit, we analyzed likelihood that both mural and septal leaflets in heart have cells labelled in the same combination of colors and compared this result to the fraction expected from random combinations (null model) estimated from the unicolor and bicolor statistics (see Figure S9). When comparing any two regions, and in particular the septal and mural leaflets, we observed a significant coincidence in the color expression of both leaflets compared to that expected from random combinations (Figure 7C). Taken together, these observations are consistent with a negative-binomial dependence of clone sizes and suggest that both leaflets may originate from a common pool of no less than 40 progenitors. This number was estimated by the number of progenitors required to cover the 2D-projected area of the leaflets, so the true number of progenitors is likely to be larger.

**Figure 7:**
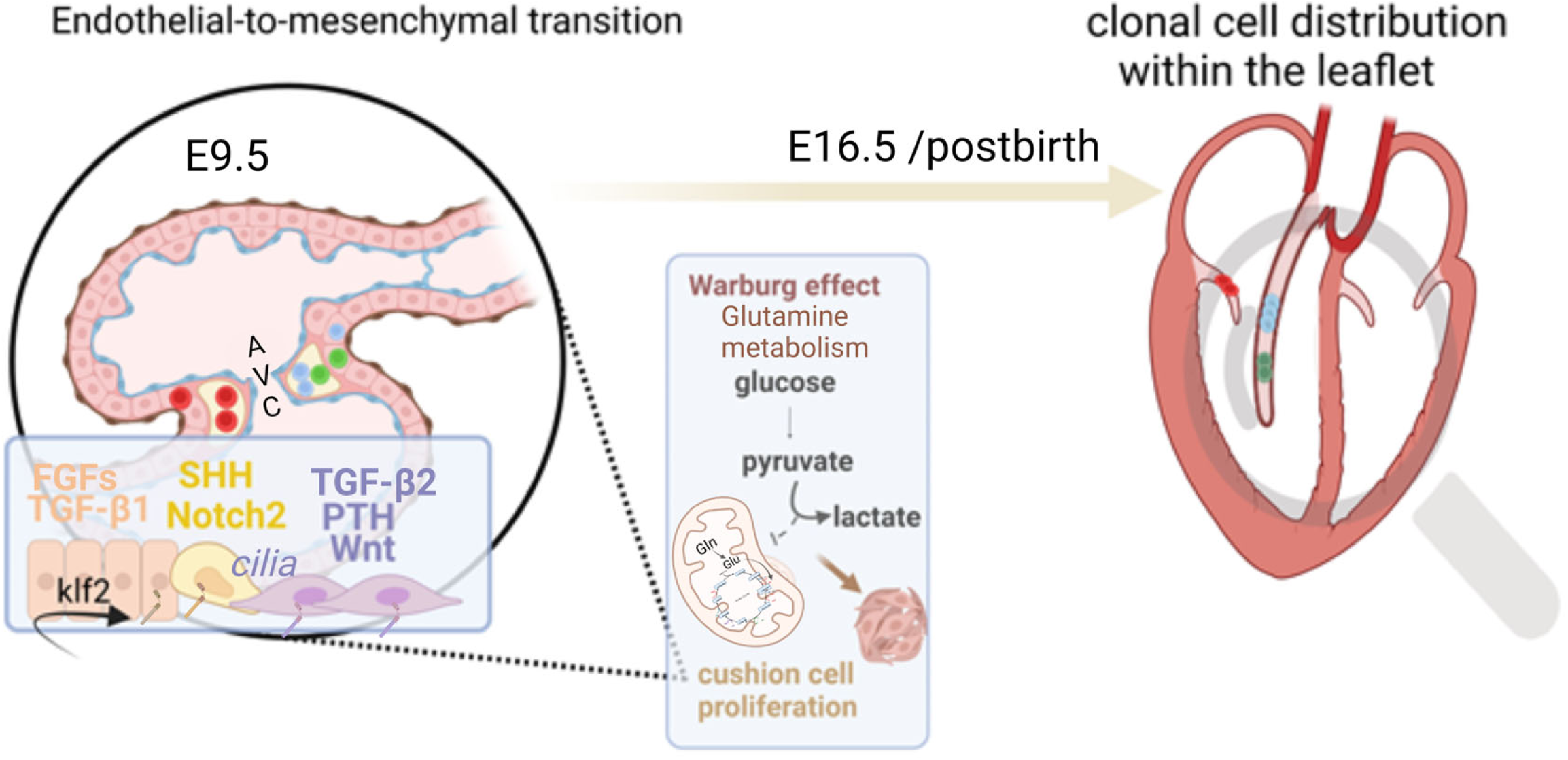
graphical summary of the formation of the mitral valve. EMT of endocardial cells at E9.5 in the atrioventricular canal (AVC) depends both upon activation of specific growth factors, hormones and transcription factors and of metabolic states of the cells. Then, specified valvular progenitors further differentiate into specific valvular endothelial (VEC) and interstitial valves (VIC). VICs distribute within the valve leaflet so they can secrete the proper extracellular matrix for a proper function of the valve

## DISCUSSION

Single-cell RNA sequencing of AVC endocardial cells and early cushion cells revealed their heterogeneity pointing to their plasticity and multipotency (Zhang et al., 2018). This further showed that a subset of endocardial cells, competent for EMT, were already primed toward a specific valvular cell type (smooth muscle cell, fibroblast, chondrogenic progenitors, endothelial cells of future endothelial layers of valve leaflet).

First, we found that a subset of endocardial cells acquired a competency to undergo EndMT. These cells expressed specific markers of EMT recapitulating the different stages of this process, as previously found in the context of cancer biology (Castaneda et al., 2022). Thus, EndMT, which allows cells to migrating out of the endocardium to form the cushions, is a cell transformation process and likely involves a finely tuned cell reprogramming process, as recapitulated in tumor cells (McFaline-Figueroa et al., 2019). Pathological EMT has been reported to be regulated by metabolism involving acetyl-CoA (Sanchez-Martinez et al., 2015) and the Warburg effect as well as by glutamine metabolism. We further found that the endocardial cells competent for EndMT turned on a specific transcriptional program encoding proteins involved in Hif1α−regulated Warburg effect and acetyl CoA metabolism such as *acly, and acss2, pfkm,ucp2, acaca and cdkn1c*.

We further uncovered that the glutamine metabolism participating in amino acid synthesis and mTorc1 pathways the latter being active during all valve formation (supplemental data 2) was likely to be very active as early as during EndMT of endocardial cells. Such an active metabolism of amino acid and protein synthesis as well as mitochondrial biogenesis is crucial for the synthesis of the ECM proteins. This transcriptional program, which likely underlies the regulation of EndMT by metabolism (Xiong et al., 2018), was turned on at the earliest step of EndMT and lasted throughout the process, as suggested by the prolonged expression of *acly* (Figure 3) as well as in the valves at E16.5 and post-birth (Figure S3).

*Ptch1, PTH1R,* and *BMPR2*, *PDGFRs* and *FGFR1* are the receptor genes to be expressed in endocardial and early *twist1*+, *prrx1,2*+ cells pointing to their role in the initiation of EndMT. Interestingly SHH signaling component early expressed during the process of EMT regulates fatty acid metabolism (Bhatia et al., 2011), while *Flt1* mainly expressed in endocardial cells prevents this metabolism rather favoring regular glycolysis (Seki et al., 2018). *PTH* was not expressed in AVC which suggests a role of maternal hormone in activation of the receptor. Our data thus suggest that only cells that specifically and in a combinatorial manner *activate PTH, Notch2, FGF, PDGF, SHH and BMP pathways* are capable of undergoing EMT while switching their metabolism toward a Warburg effect (Liberti and Locasale, 2016), glutamine metabolism and fatty acid synthesis. The glutamine availability and in turn its metabolism is likely to be sensed (Steidl et al., 2023) by a subset of endocardial and cushion ciliated cells (Burns et al., 2019; Van der Heiden et al., 2006). This cilia-regulated metabolism and specific EndMT competency of a subset of endocardial cells likely explain that EndMT occurs only in the local atrioventricular canal which features a change in blood flow and thus in stretch forces sensed by endocardial cells when compared to endocardial cells lining both chambers (primitive ventricle and atrium).

Expression of *hand2* and *msx1* are all required for endocardial cells to undergo EndMT and to be committed toward a valvular cell. These endocardial cells further loss *cldn5*, a claudin and *gja4*, a connexin that limits endothelial cell motility (Aasen et al., 2019; Yang et al., 2021) and turn on expression of *cldn6* and *cldn7* as well as *gja1* involved in cancer metastasis (Aasen et al., 2019; Tabaries and Siegel, 2017) and in turn in migratory mesenchymal cells. Collectively, our data explain why only a restricted population of cells follows this EndMT program.

As recently reported as a general concept, endocardial cell state heterogeneity makes cells either low or high responder to growth factors, cell stretch and downstream signaling to ensure specific and context-dependent cellular decision in a multicellular setting (Kramer et al., 2022).

The EndMT transcriptional and cell metabolic reprograming events are then followed in mesenchymal cells by a genetic program specific to future valvular cell types. The latter is designed to allow cells to acquire a capability to secrete specific extracellular matrix proteins (ECM) (Figure S2). We found that, at E16.5, valvular cells expressed genes encoding ECM proteins found in layers of the valve leaflets, while a subset remained endothelial to form the surrounding layer of the leaflets. The process was more advanced at the neonatal stage, with the mitral valve featuring two different types of VICs expressing collagen genes as well as proteoglycan genes. Interestingly the VECs could be discriminated between parietal and septal valve leaflets depending upon the presence or absence, respectively, of cells expressing *wt1*. Wt1 lineage tracing indeed was reported to contribute to VICs as well as VECs of the parietal mitral leaflet (Wessels et al., 2012) (Figure S1).

This suggests that some cells of (pro)epicardial origin expressing Tie2 at the time of induction of the recombinase retained expression of *wt1* even after migration into the valve leaflet. Interestingly VEC could still undergo EMT to provide after birth the leaflet with more VICs (Figure 5F). Hemogenic cells from endocardium also contributed to neonatal VIC likely through generation of macrophages in the endocardium to remodel the valve (Shigeta et al., 2019). Altogether, these findings are in agreement with the long process of maturation of valves in early adult (Hulin et al., 2019) in which macrophages from bone marrow also contribute to (Kim et al., 2021).

The VICs of both parietal and septal leaflets likely originated from the same AVC cells cluster (Figure 5F) which was in agreement with both the single cell transcriptomics data processed within a trajectory inference (Figure 4D,5) and the clonal analysis that also points to the same pool of progenitors shared by both valve leaflets (Figure 6). Clonal analysis based on 2D leaflet areas predicted that at least 40 progenitors were required to cover the area of the septal leaflet and around 20 for the parietal leaflet. Considering the 3D morphology of the leaflets, this figure is broadly consistent with the total number of AVC cells that contribute to the E16.5 Mitral leaflets based on single-cell data (obtained by taking the number of cells in each contributing cluster (Figure 5F) and dividing by the number of embryos, which led to a figure of around 140 AVC progenitors). This implied that the total clone size results from a growth process of differentiating progenitors that expand before terminally differentiating in VICs. Assuming a number of cells in a mitral valve leaflet of 400 in septal and 150 in mural leaflets at birth (Moore et al., 2021), this suggests that progenitor cells should undergo about three rounds of division on average.

Another interesting feature in the behavior of cells is their distribution along the long axis of the leaflet (Figure S6). Such a cell behavior could explain the deposition of the ECM as layers, a characteristic of these valvular leaflets.

Altogether, our single-cell RNA-sequencing data combined with the clonal analysis revealed that the process of embryonic mitral valve formation starts early during development when AVC endocardial cells distribute between EMT-competent valve progenitors cells and endothelial/endocardial cells that will participate in lining the chambers during their growth, respectively. The EMT process is restricted to cells sharing specific cell identity (i.e., unique transcriptomic profile, competency to respond to specific agonists, to react to fine-tuned stretch and ability to undergo a metabolic cell reprogramming event) (Figure 7). Restriction of such a cell population with the previous criteria may prevent any overgrowth of the cushions and lead to a well-tuned size of the leaflets, mandatory to ensure a perfect closure of the future valve.

We found also that cells specifically contribute to each anterior or posterior cushion and later to specific mural or septal leaflet. Other cells from a different embryonic origin (Farrar et al., 2021) migrate later into these respective valve leaflets to complete the whole process of leaflets formation. The 2D distribution of the cell clones and their transcriptomic profile at E16.5 including ECM genes (Figure 4, Figure S2) indicate that the cell clones are specified toward the spongiosa, the fibrosa or the atrialis. The clones likely contribute to the formation of specific layer of the valve. Dysregulation of any of these processes may lead to a pathological valve.

## Acknowledgments

This study was funded by the Leducq Foundation (MP, MITRAL network of excellence). We are also grateful to the Leducq Foundation for generously awarding us for cell imaging facility (MP “Equipement de Recherche et Plateformes Technologiques” (ERPT)).

This study was also partly supported by Agence Nationale de la Recherche (contracts ANR-11-EQPX-0029 Morphoscope2 and ANR-10-INBS-04 France BioImaging, to EB and IHU FOReSIGHT, ANR-18-AHU-01 and ANR-19-CE16-0019 to JL).

We thank HALIODX (Marseille) for running the single-cell RNA-seq experiments Figure 1A and 7 were designed using Biorender.

## Methods

### Mouse lines

*Rosa26–tdTomato and Tie2cre* mice were obtained from Jackson laboratory (B6;129S6-*Gt (ROSA)26Sortm14(CAG-tdTomato)Hze and B6.Cg-Tg(Tek-cre)12Flv/J, respectively)*. The WT1*^CreERT2^*mouse and *CAG-CreER^TM^* mice (Guo et al., 2002) were kindly provided by Sylvia Evans (UCSD) and Joshua R. Sanes (Harvard University), respectively. The *Cytbow* mouse line broadly express a *CAG-Cytbow* transgene designed on the model of Loulier et al(Loulier et al., 2014), modified to stochastically express mTurquoise2, mEYFP, or tdTomato from the broadly active *CAG* promoter following Cre recombination (this line will be described elsewhere).

Mice were kept under standardized conditions (22–24°C temperature; 50%±10% humidity) on a 12 h light/12 h dark cycle in an enriched environment (kraft paper enrichment). Food and tap water were provided ad libitum. The parental strains were bred on a C57BL/6 background.

*CAGCreER^TM+/-^* mice were crossed with the transgenic mice *cytbow*. To obtain embryos of the desired stages (E9.5, E16.5), mating females were verified for the presence of vaginal plug every morning. The day of the presence of a vaginal plug was considered as embryonic day E0.5 at noon. Pregnant females were euthanized at the desired stage of gestation.

Tamoxifen (Sigma-Aldrich, France; 20 μg/g) was administrated with corn oil by oral gavage at embryonic day E8.5. Embryos were collected at E9.5 and E16.5, fixed overnight with 4% paraformaldehyde.

### Preparation of frozen sections

Embryos at E9.5 and hearts at E16.5 were prepared for immunostainings. Samples were embedded in Optimum Cutting Temperature (O.C.T) compound (Tissue-Tek) for cryoprotection. Embryos at E9.5 and E16.5 were dissected in ice-cold PBS and fixed for 2hrs or overnight at 4°C in 4% PFA in PBS. Embryos were then transferred to 6% sucrose/PBS (w/v), 12.5% sucrose/PBS and 25% sucrose/PBS (1hr each) with gentle shaking at 4°C. Then the embryos were transferred into embedding molds, and frozen in 50:50 (v/v) O.C.T/50% sucrose. Samples were stored at -80°C and sectioned (10μm, 20μm, 30μm) using a Leica Microm HM 500 OM Cryostat (-20°C). Frozen sections were mounted on superfrost plus slides and stored at -80°C before use.

### Immunofluorescence analysis

Immunofluorescence was performed on frozen embryonic heart sections (Leica cryostat-10 μm). Sections were fixed with PFA for 10 minutes, washed with PBS 5 minutes, then blocked in PBS with 0.5% donkey serum, 5% BAS and 0.1% tween 20. Sections were stained with the following antibodies: anti-CD31 (BD550274 Purified Rat anti-mouse CD31; 1/100), anti-sarcomeric alpha actinin (mouse, Sigma-Aldrich A7811; 1/200) anti Twist1 (mouse 1/300 Novusbio NBP2), anti-ACLY (37364SS ACLY rabbit 1/200 Novusbio NBP2 – 67509), anti-Msx1 (rabbit 1/300 Novusbio NBP2 – 57021), anti-VECAM-1 (rabbit 1/250 Abcam ab-134047), anti-Integrin alpha V (rabbit 1/500 Abcam ab-179475) anti-Integrin beta 3 (rabbit 1/500 Abcam ab-119992). Nuclear staining was performed with DAPI (1/10000).

Images were acquired using a confocal LSM 800 Zeiss microscope equipped with an airryscan of 32 detectors. Light was provided by a laser module 405/488,561 and 640 nm wavelengths, respectively.

All images were acquired using the ZEN ZEISS software. Then some images were deconvoluted using Autoquant and reconstructed in 3D using Imaris software (IMARIS). All samples were mounted in Fluoromount™ (Cliniscience, France)

In all figures, tdTomato/mCherry, mEYFP, and mCerulean/mTurquoise2 are rendered in red, yellow, and blue, respectively.

### Two-photon imaging of whole embryonic hearts

Large-scale color two-photon microscopy of whole embryonic hearts was performed on a custom-built imaging platform(Abdeladim et al., 2019) combining trichromatic two-photon excitation through wavelength mixing(Loulier et al., 2014) and 2D mosaicking of XYZ image series. Hearts were clarified following the CUBIC procedure with a modified final solution(Susaki et al., 2014) and mounted between two coverslips and rigid spacers. Simultaneous excitation of CFP, YFP and tomato was performed by mixing 850 and 1150 nm pulses focused by a multiphoton water-immersion objective equipped with a correction collar (25×, 1.05NA, XLPLN25XWMP2, Olympus, Japan). Detection was performed on three independent detectors equipped with appropriate filters (Semrock FF01-475/64-25, Semrock FF01-542/27-25, Semrock FF01-607/70-25). The correction collar was set to obtain optimal contrast at a depth corresponding to the middle of the sample. A total volume of 1580 × 1580 × 300 µm3 was recorded in the form of 4× 4 XYZ stacks of dimensions 420 × 420 × 300 µm3 with voxel size 0.42 × 0.42 × 1.5 µm3. The volume was reconstructed using ** Fiji / bigstitcher** (Horl et al., 2019) and imported in Imaris (Bitplane, Switzerland) for visualization.

### Assessment of the recombination of the *cytbow* reporter by PCR

Genomic DNA was extracted by DNA precipitation in NAOH solution (50 mM) from the tail of mice, wild type mouse and adult *cytbow* mouse as a control. Tails of embryos at E16.5 *CAG-CreER^TM+/-^/cytbow* of the whole litters were used to genotype embryonic mice with specific primers.

### Mouse Genotyping Primers

CYT forward 5’-CTGTTCCTGTACGGCATGGA-3’ and reverse 5’- TTTCAGGTTCAGGGGGAGGT-3’; CAG^CreERT2^ 5’GCTAACCATGTTCATGCCTTC-3’; 5’- *AGGCAAATTTTGGTGTACGG-3’; ROSA26^tdTomato^; Wild type, forward: 5’- AAGGGAGCTGCAGTGGAGTA-3’ and reverse:5-’CTGTTCCTGTACGGCATGG-3’, Mutant, forward:5-’CCGAAAATCTGTGGGAAGTC-3’ and reverse:5’- GGCATTAAAGCAGCGTATCC-3’; Tie-2^Cre^* forward *5’- CGCATAACCAGTGAAACAGCATTG-3’ and reverse 5’- CCCTGTGCTCAGACAGAAATGAGA-3’; WT1^CreERT2^ forward 5’- TGAAACAGGGGCAATGGTGCG-3’ and reverse 5’-CGGAATAGAGTATGGGGGGCTCAG- 3’*.

### Single-cell RNA sequencing

Endocardial cells of the atrio-ventricular canal derived from Tie2*^Cre^*/Tomato positive embryos at E9.5 (18-22 somites) and mitral valve leaflets were collected at E16.5 and P1 pups, dissociated for 10 min with TrypLE^TM^ express enzyme (Gibco; Catalog number 12604013) in the presence of Y-27632 Rock inhibitor (stem cell technologies) into single cells. After sorting the tomato positive cells with FACS, 10x genomics chromium platform was used to capture and barcode cells in order to generate single-cell Gel Beads-in emulsion (GEMs), following the manufacturer’s protocol. Thus, using the reverse transcription master mix, cell suspensions of 2 samples of AVC, 1 sample of E16,5,and 2 samples of P0 were loaded onto the 10x Genomics Single Cell 3’ Chip. Single cells were then partitioned into GEMs along with gel beads coated with oligonucleotides. Reverse transcription of cDNAs was followed by amplification, and a library was constructed using the Single cell 3’ reagent Kit for all the samples, followed by sequencing of all libraries wit

Next-seq Illumina sequencer. 10X Genomics pipeline Cell Ranger v1.2.0 was used for sample demultiplexing, barcode processing, and UMI counting. Libraries obtained from Illumina Sequencer were demultiplexed into reads in FASTQ format using the “cell ranger mkfastq” pipeline. In order to generate a gene-barcode matrix for each library, cellranger count pipeline was used. All the reads were aligned to the *mus musculus* reference genome mm10.

Cell ranger pipeline was used to normalize all the libraries to the same sequencing depth. A first analysis of the data was performed with cell ranger and loop cell 10xGenomics software’s. C loop clustered the cells using genes that are differentially expressed comparing to all other cells in other clusters with a threshold of Log2 > or equal to 2. A deep and secondary analysis was performed using the SCDE package and SCShinyhub(Jagla et al., 2021). Low UMI cells were excluded before clustering. We used: for E9.5 samples: sample: 1200 cells/ median gene per cell: 4617. For E16.5 sample: 452 cells/ median gene per cell: 700/; For P0 2 samples were used for a total of 854 cells/ median gene per cell 900.

### Clonality assessment and clonal grouping

Cell labeling followed tamoxifen administration at E8.5, and samples were collected at E16.5. At collection, labelled cells had expanded into clones that appeared to be fragmented into separate clusters of cells within a leaflet. To infer whether a group of cell clusters belonged to a single clone or were the result of merger of two or more clones, we followed a biostatistical approach(Lescroart et al., 2014). In this approach, one considers the probability *p* that a given cell was induced, as shown in Figure S4. The probability *P*(*m*), of observing *m* induction events in a given leaflet can be approximated by a Poisson distribution: 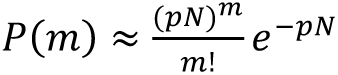 **(1)**, where *pN* is the overall induction probability, and N is the number of cells expressing the promoter. If clone fragmentation events are considered to be random and statistically uncorrelated (i.e., Poissonian), the probability *R*(*k*) that a given clone will end up partitioned into *k* fragments is: 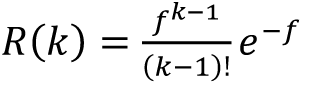 **(2)**, where *f* corresponds to the expected number of fragments of a clone derived from a single cell at the time of measurement (E16.5). As argued in lescroart et al. (Lescroart et al., 2014), the degree of fragmentation *f* could depend on the total size of the clone. To assess this, we measured the relation between total clone size and number of fragments in unicolor hearts (extended data Fig. 6). Although the number of measurements restricted to leaflets labelled with a single color was low, there was no clear dependence between these two variables. Note that, here, we considered clones in separate leaflets in the same heart to be independent. Noting that the number of fragmentation events is given by the difference *k-m* (*k≥m*) between the number of fragments *k* and the number of induced cells *m*, then the conditional probability *S* (*k|m*) of finding a total of *k* labeled fragments of a single color at day E16.5, given that *m* cells of that same color were induced at day E8.5 is given by 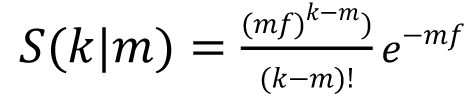 **(3)**. Combining equations **(1)** and **(3)**, we obtained the joint probability *J* (*k|m*) of finding a leaflet with initially *m* induced cells giving rise to *k* fragments of a given color is given by

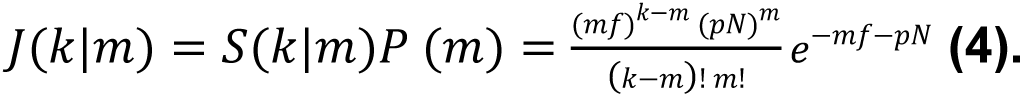

Taking into account the ensemble leaflets in which there is at least one induced cell (*m >* 0), we defined the joint distribution of labeled hearts 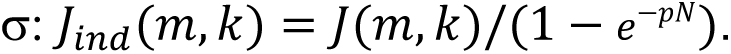. Then the distribution for the number of fragments in labeled hearts is given by:

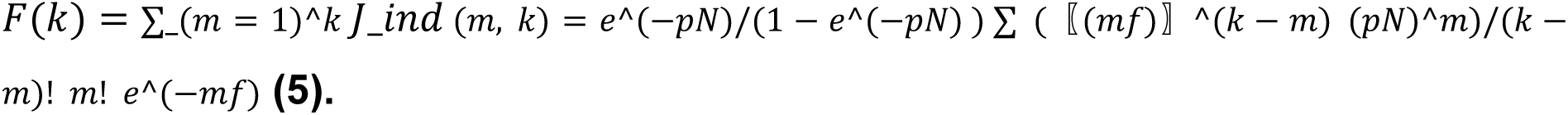

To assess the clonality of our lineage tracing data, we used the experimental quantification of the number of fragments *k* and Eq. **(5)**. We first estimated the overall induction frequency *pN* and average fragmentation rate of clone *f* by combining the data from both the mural and septal leaflets. Given the spatial separation between the mural and septal leaflets, we considered the cell clusters from separate leaflets to be independent from each other. Thus, if in a given sample we observed e.g., 4 red fragments in the septal and 3 red fragments in the mural leaflet, we considered these to be two clonal induction events, one with 4 and the other with 3 fragments. We analyzed the number of fragments in both leaflets of the mitral valve of 40 hearts, from which we obtained fragments for 171 clone candidates. We analyzed the distribution of the number of fragments (extended data Fig. 7) and fitted the theoretical curve (**5**), from which we obtained an overall induction frequency *pN* = 0.5 (0.3, 95% C.I.) (**2**), and a fragmentation rate *f* = 1.4 (0.3) (**3**). This implied that clones with 4 or less fragments were likely to be of clonal origin, when the data for both the mural and septal leaflets was pooled together.

Even though the induction frequency *pN=0.5* can be assumed to be equal for both leaflets, this is not necessarily the case for the fragmentation rate *f*.

To account for possible differences in the fragmentation in the mural and septal leaflets, we analyzed the distribution of number of fragments *F*(*k*) of both leaflets independently (Figure 5) for fixed induction frequency *pN* = 0.5, as obtained from the combined data (Figure S7). In the case of the septal leaflet, the threshold for clonality was found to be *k*=4, meaning that groups of four clusters or less were considered clonal, while this threshold was lowered to *k*=3 for the mural leaflet. The difference originates from the lower fragmentation *f* in the case of the smaller mural leaflet.

## Notes

### Competing Interest Statement

The authors have declared no competing interest.

### Summary of Updates

A single cell transcriptomic and clonal analysis depicts valvulogenesis new data have been added

