## supplemental figures for "Birth, cell fate and behavior of progenitors at the origin of the cardiac mitral valve"

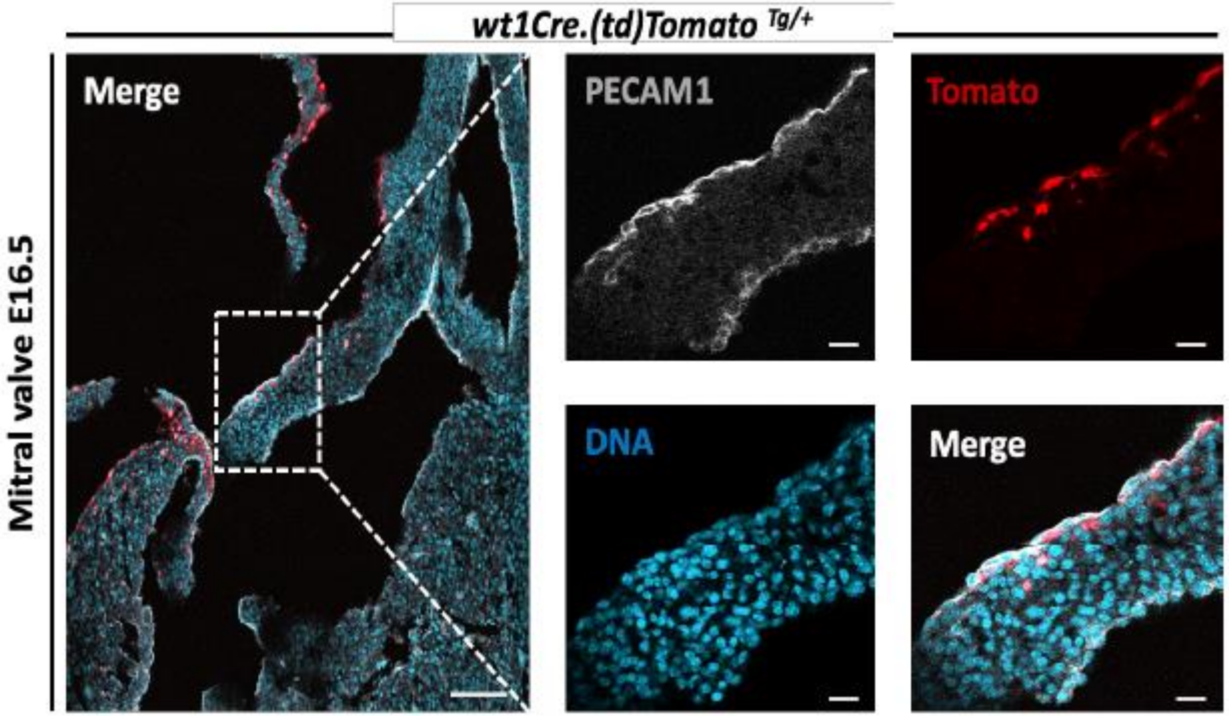


**FIGURE S1** Lineage tracing of epicardial cells. Wt1$^{creert2}$ bred with Rosa26 $^{tdTomato}$ mice. The cre was induced by tamoxifen gavage to the mother at E8.5. The embryonic hearts were collected at E16.5 and the mitral valve imaged. The parietal leaflet is magnified in the right images

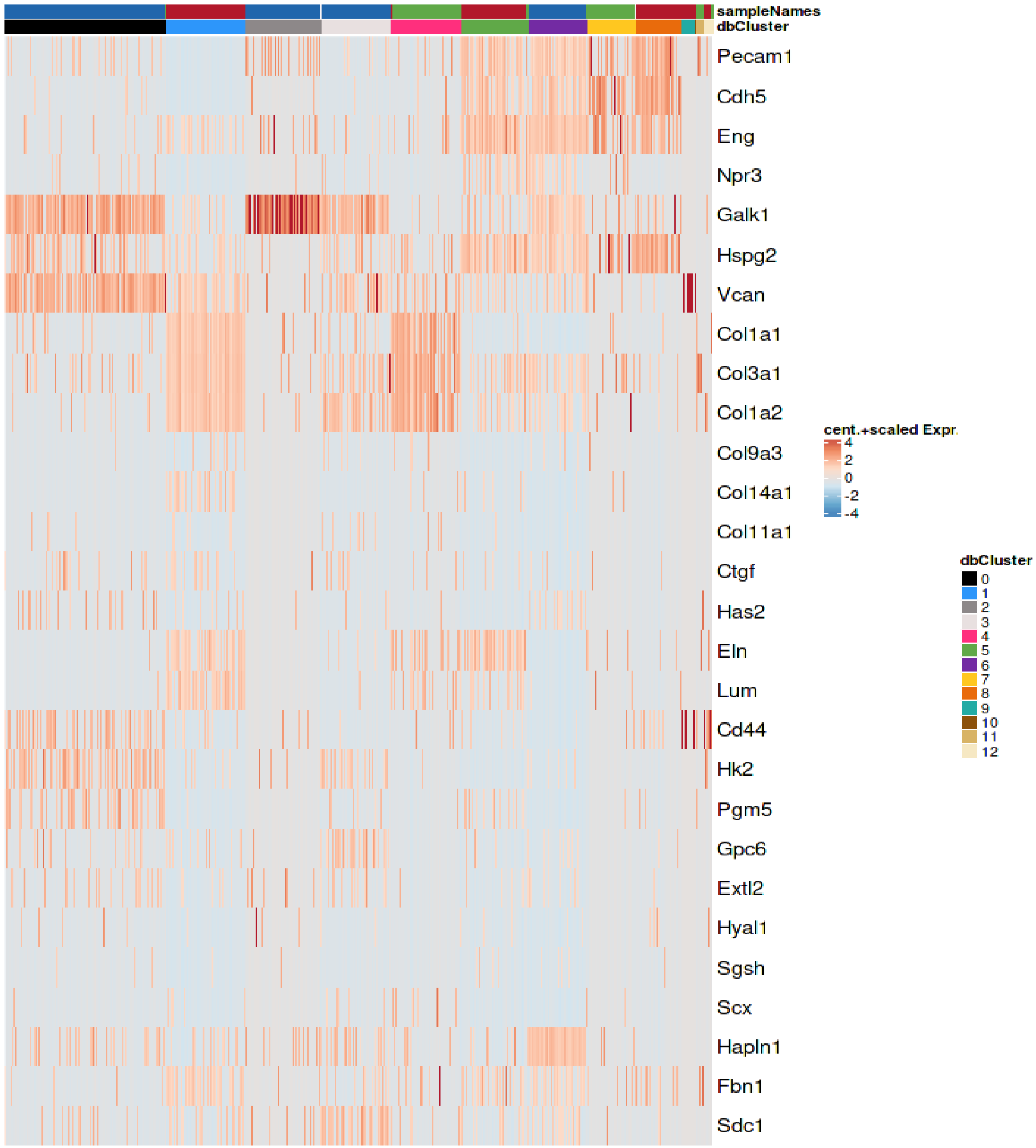


**FIGURE S2:** Heatmap of cells revealing expression of genes involved in extracellular matrix ( ECM) production

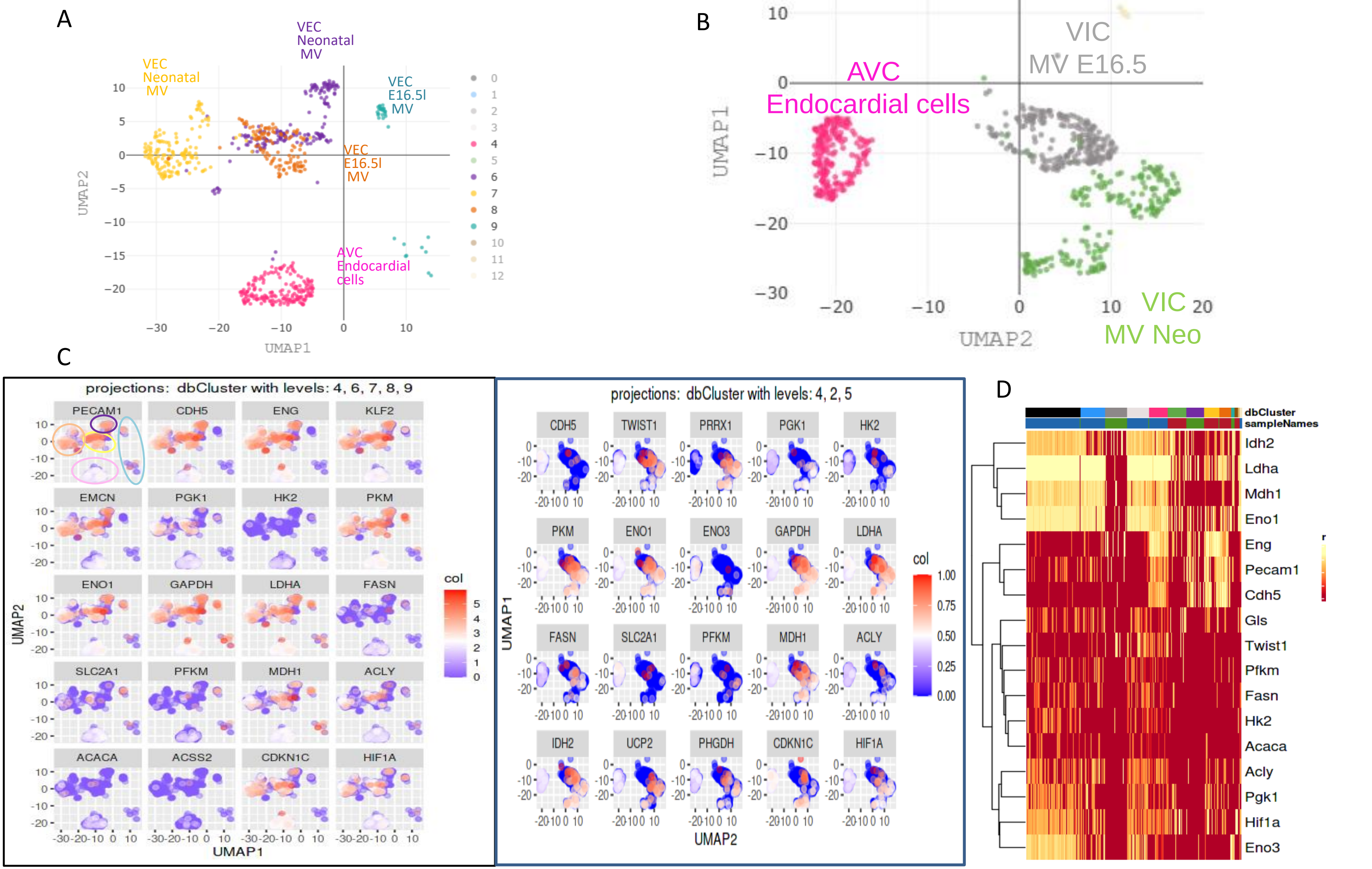


**FIGURE S3:** (A) UMAP of endocardial cells and VECs of E16.5 and neonatal mitral valve (B) UMAP of endocardial cells and VICs of E16.5 and neonatal mitral valve (C) respective panel plots of genes involved in Warburg effect in endocardial cells ( cluster 4) and VECs clusters 6,7, 8, 9 ) or VICs (clusters 2, 5) of E16.5 and neonatal mitral valve (D) heatmap of genes involved in Warburg effect

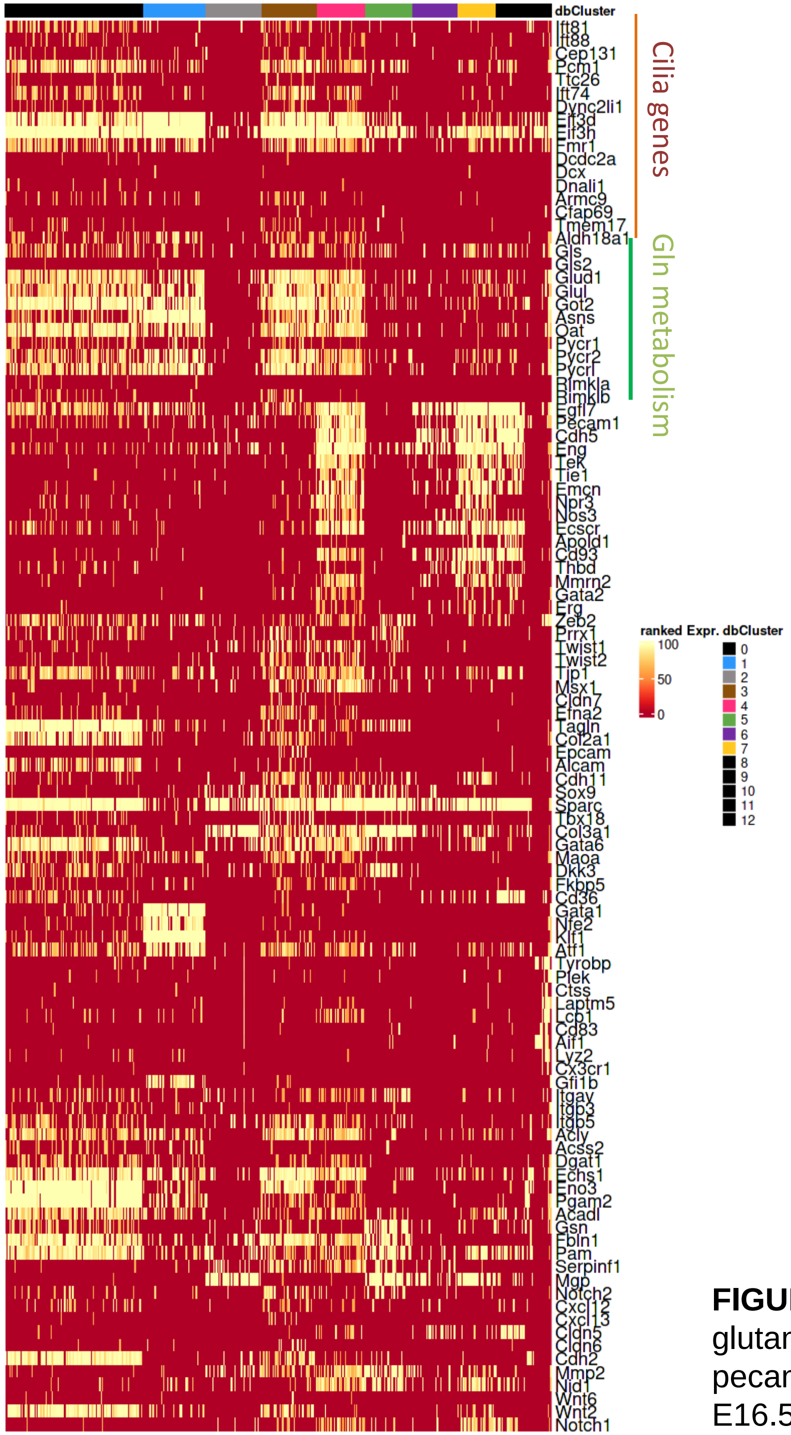


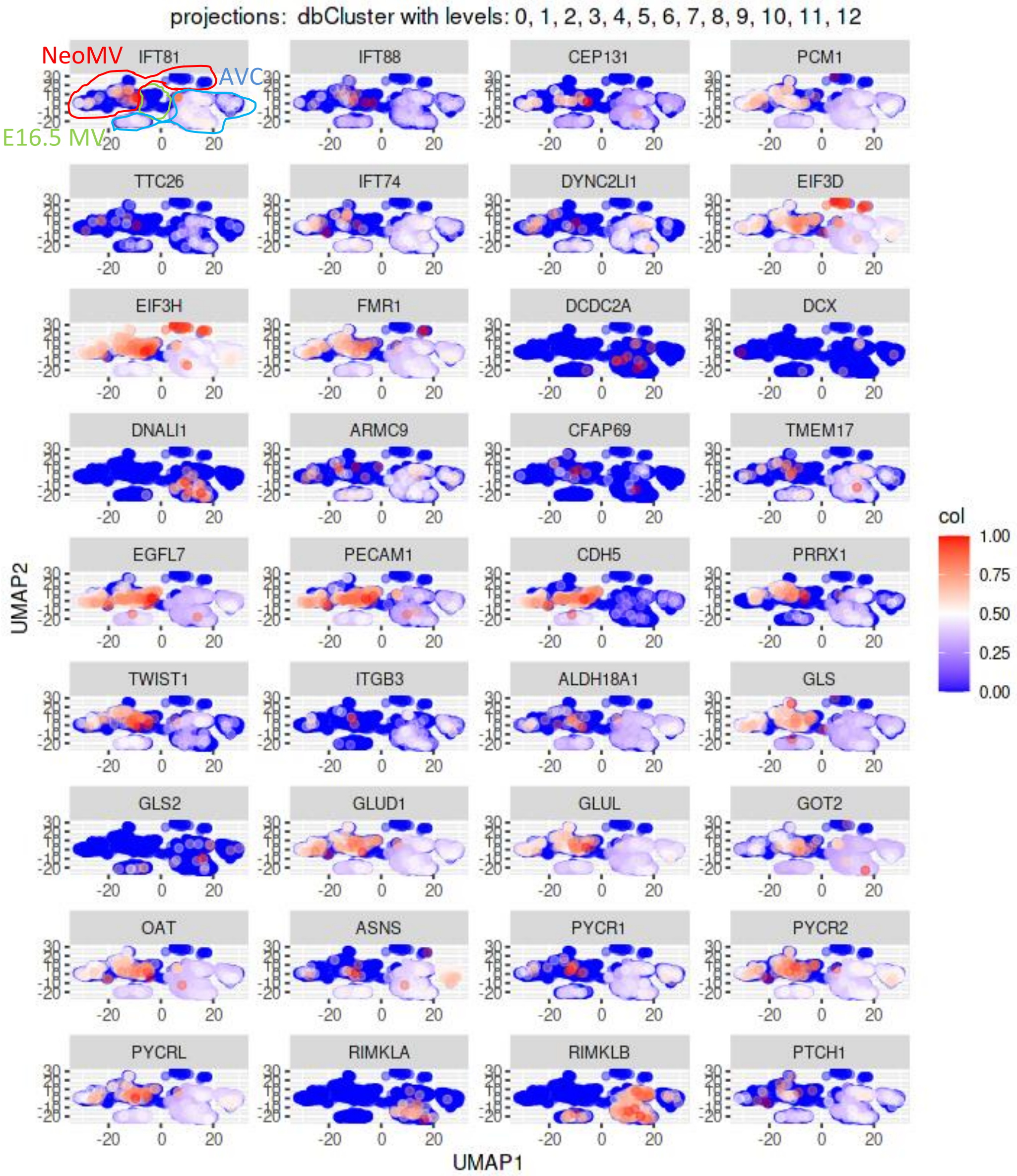


**FIGURE S4** expression pattern of genes involved in cilia formation and glutamine metabolism in the 3 developmental stages of valve formation; pecam1, egfl7 and cdh5 mark both endocardial cells in AVC and VECs in E16.5 and neonatal mitral valves

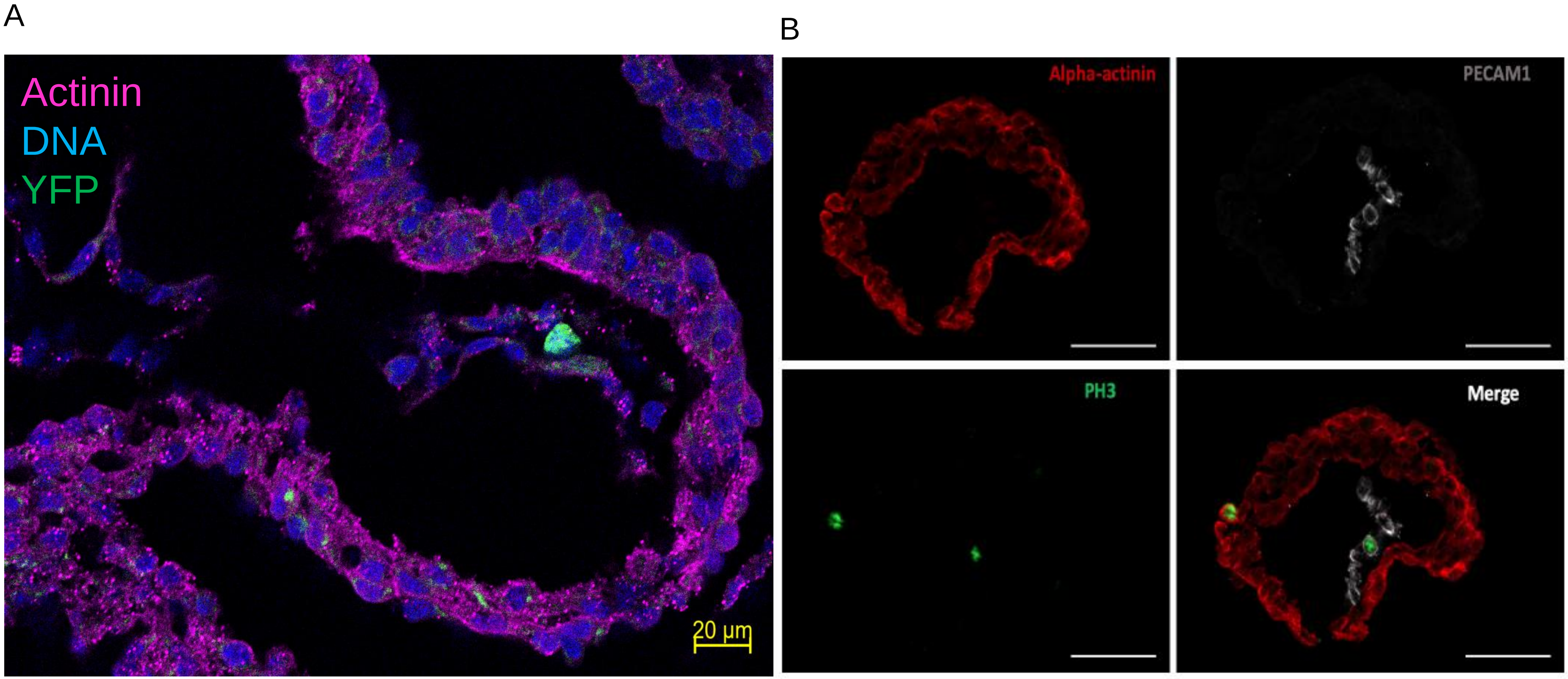


**FIGURE S5:** (A) E9.5 embryonic heart from CAG$^{creERT2}$ x Rosa26$^{TCY}$ breeding mice revealing a single YFP+ cell in the AVC.The female was given a low dose of tamoxifen (20 µg/g) at E8.5 of gestation. (B) E9.5 heart section stained with a anti-PH3 antibody reveals a low cell division rate in the endocardium

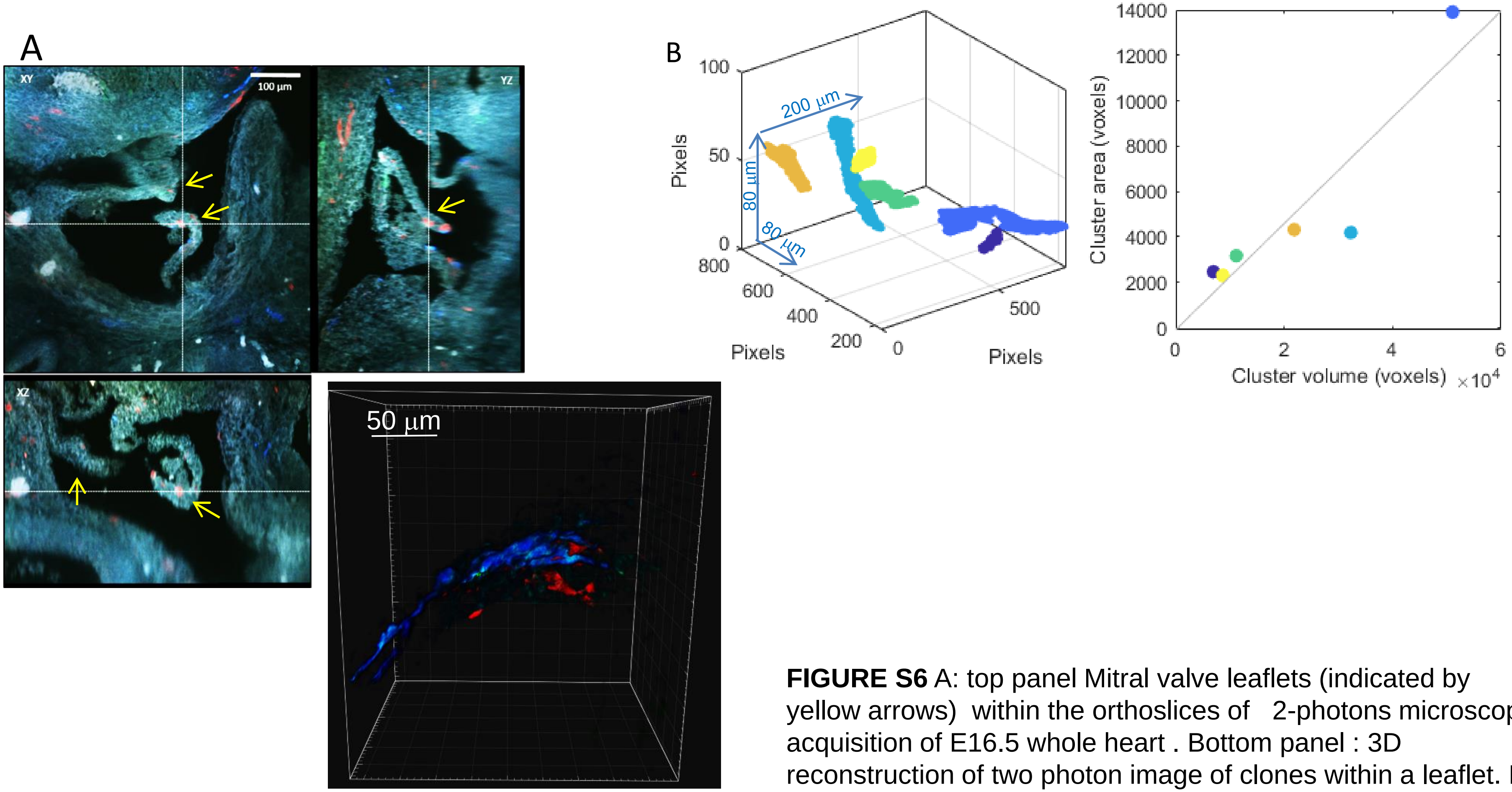


**FIGURE S6** A: top panel Mitral valve leaflets (indicated by yellow arrows) within the orthoslices of 2-photons microscopy acquisition of E16.5 whole heart . Bottom panel : 3D reconstruction of two photon image of clones within a leaflet. B: correlation between the 2D-projected area of cell clusters and their 3d volume

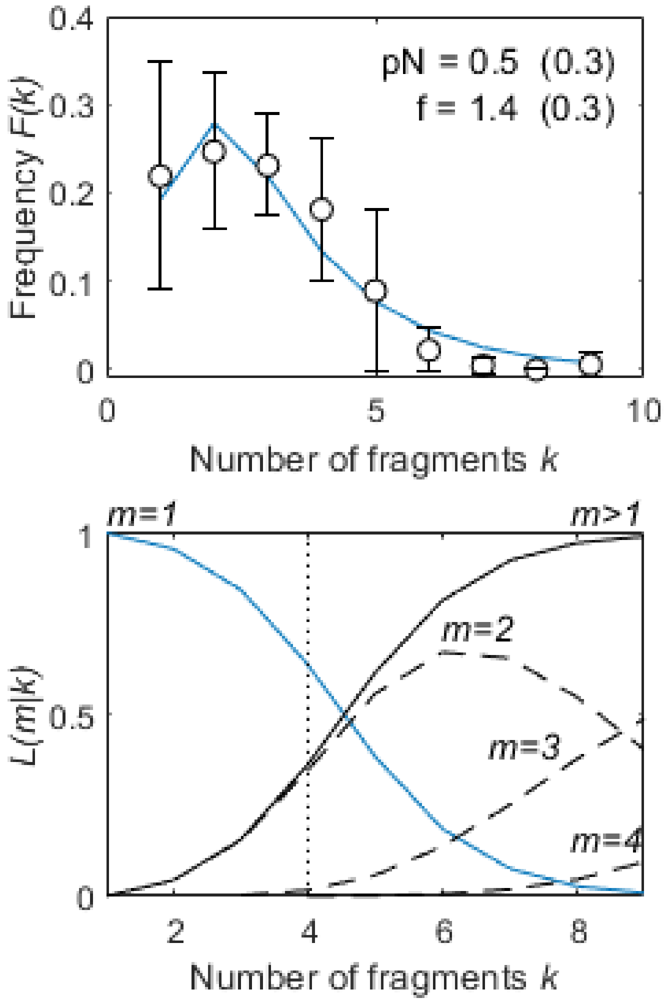


**FIGURE S7:** Overall induction frequency versus the clone fragmentation rate f

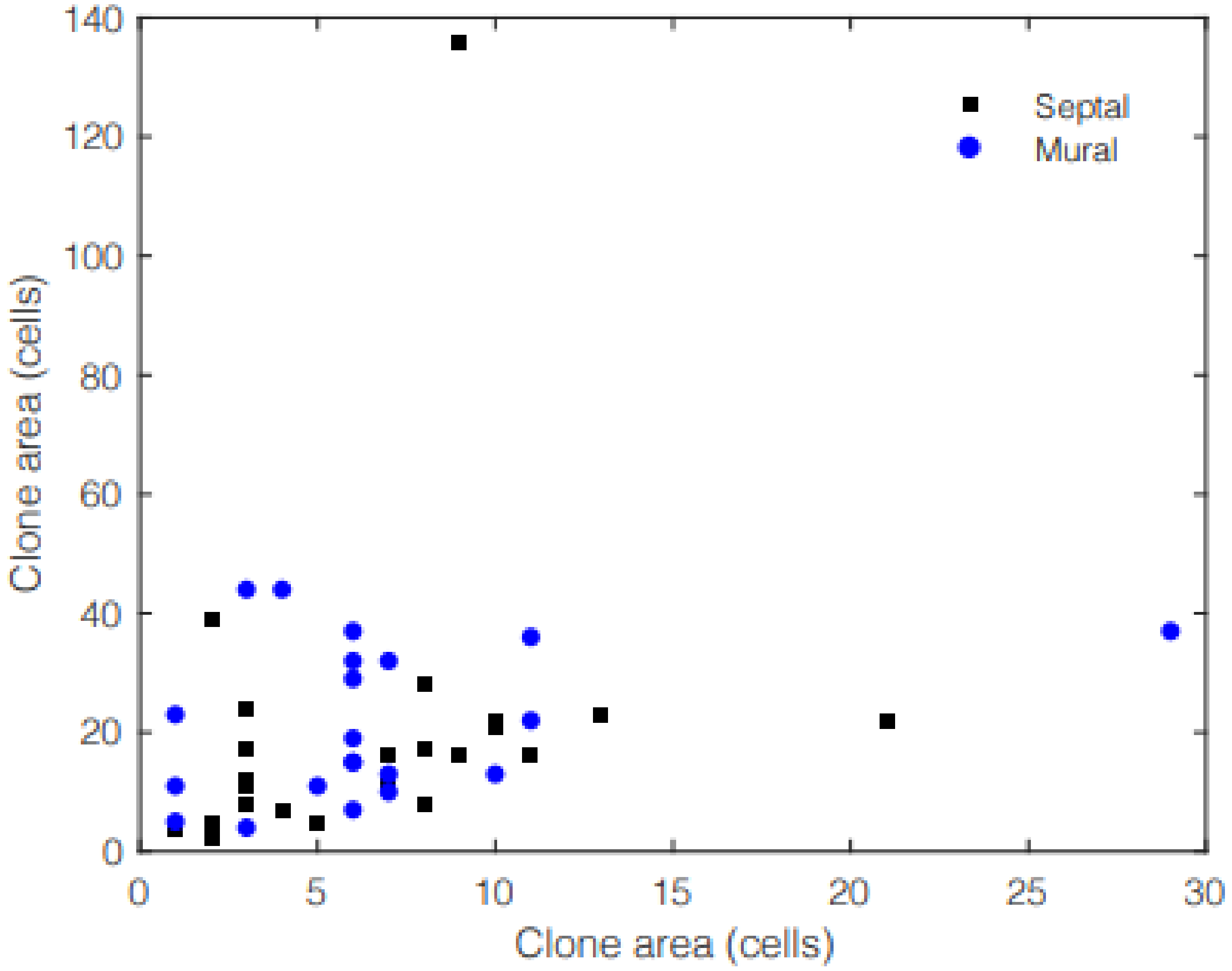


**FIGURE S8**: correlation of the sizes of clones of different colors in the same leaflet

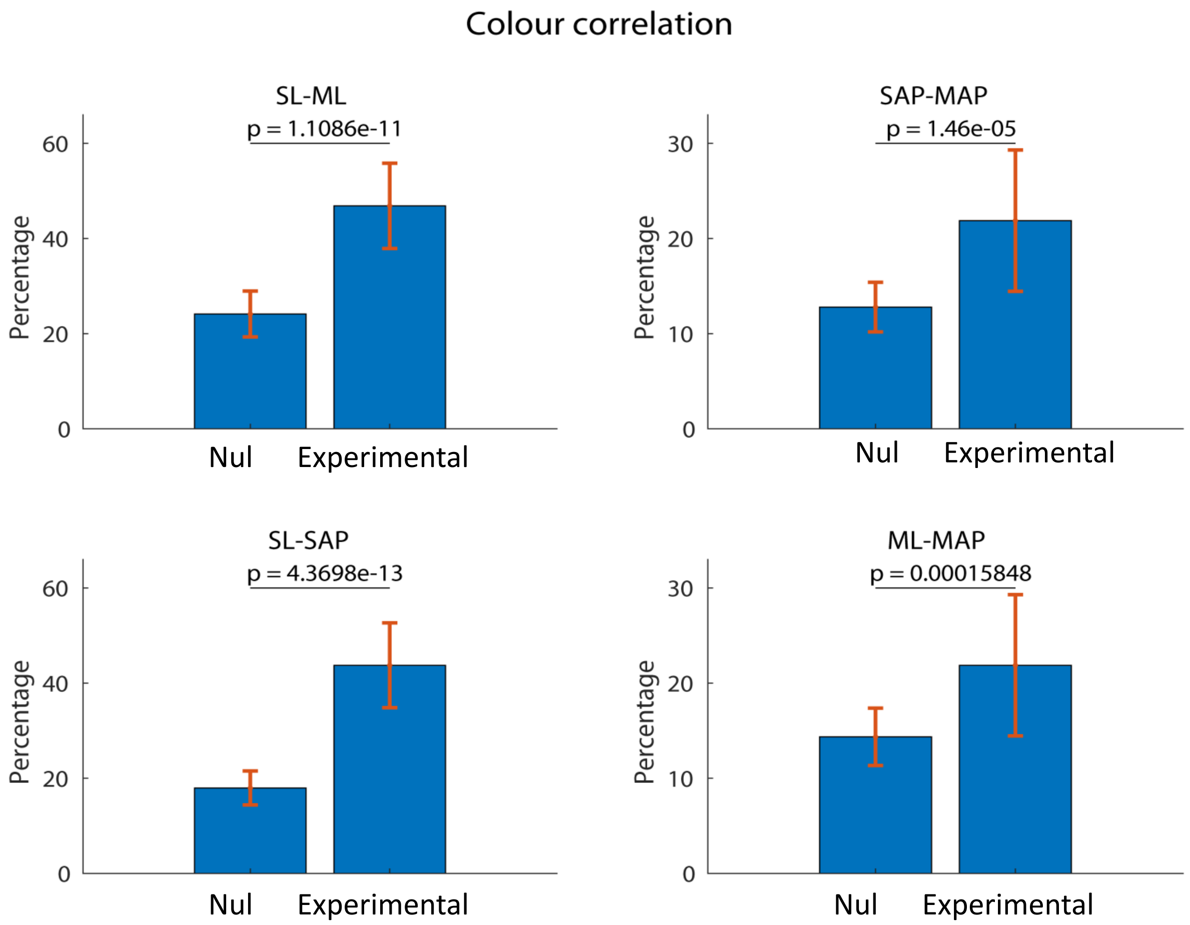


**FIGURE S9:** comparison of the fraction expected from random combinations (null model) with estimation (experimental) from the unicolor and bicolor statistics.
